# Neural and behavioral correlates of sensorimotor synchronization to a subdivided beat: Evidence for a binary grouping advantage and musical training effects

**DOI:** 10.1101/2021.02.15.431304

**Authors:** A. Celma-Miralles, B.A. Kleber, J.M. Toro, P. Vuust

## Abstract

Motor synchronization is easiest when the metronome beats are subdivided into equal intervals. Here we combine electroencephalography (EEG) frequency-tagging and mismatch responses to examine how neural synchronization and predictive processing contribute to sensorimotor synchronization across four beat subdivisions: duplets (1:2), triplets (1:3), quadruplets (1:4), and quintuplets (1:5). We recorded EEG in musicians and non-musicians during a finger-tapping paradigm combining passive listening and synchronized tapping. Each trial had four sections: an isochronous beat, one of the four beat subdivisions, the beat again and the same beat subdivision. Only in the fourth section participants had to tap the beat while listening to the subdivision. We found that musicians outperformed non-musicians in the tapping task and showed stronger neural activity synchronized to the beat frequency and its harmonics for duplets, triplets and quadruplets. Importantly, non-musicians showed better tapping performance and stronger neural responses specifically for duplets only, indicating a sensorimotor synchronization advantage for the simplest binary subdivision. Mismatch responses to unexpected sound onsets likewise converged on the same pattern: listening to the first subdivision onset elicited smaller prediction errors for duplets in both groups, suggesting that binary subdivision better aligned with listeners’ metrical expectations. Together, these findings show a correspondence between behavior and neural processes underlying the subdivision levels of the beat and highlight sensorimotor advantages for simple integer ratios that can be modulated by training in music. Both neural synchronization and predictive processes seem to jointly support this sensorimotor synchronization advantage for binary subdivisions.

## 1. Introduction

Sensorimotor synchronization (SMS) refers to the ability of the human brain to synchronize its motor actions with external rhythmic stimuli, such as tapping one’s feet while listening to music. This process involves generating temporal expectations by identifying a prominent beat within a metrical hierarchy (Large, 2008; Vuust and Witek, 2014), which is also reflected in the hierarchical organization of coordinated movement (Mårup et al., 2022). Rhythmic stimuli are perceived relative to an evenly paced pulse from which listeners infer a metrical hierarchy comprising faster (i.e., subdivisions) and slower temporal levels that provide reference points and give rise to patterns of metrical accentuation (London 2012; Honing 2012). SMS supports many human activities, including music performance, dance, and sports (Solberg & Jensenius, 2019; Damm et al., 2020), and has become an important component of movement rehabilitation (Nombela et al., 2013; Schaefer 2014; Bek et al. 2020). However, despite its broad relevance, the neural mechanisms by which beat subdivisions enhance SMS remain poorly understood, raising the question of how the brain represents beat subdivisions across different temporal scales and how musical training shapes this process.

According to predictive processing accounts, the brain represents rhythmic events across multiple temporal scales, continuously comparing predictions with incoming sensory information to minimize prediction errors and guide future actions (Koelsch et al., 2019, Vuust et al., 2022; Friston, 2010). Because successful SMS requires anticipating future beat events rather than merely reacting to them, finger-tapping paradigms provide an effective tool for studying predictive timing by measuring accuracy and variability across taps (e.g. Iversen and Patel, 2008; Dalla Bella et al., 2017). These paradigms have revealed two major effects of subdivisions on temporal perception and synchronization: (i) the filled duration illusion (Buffardi, 1971; Thomas and Brown, 1974; Wearden et al., 2007) and (ii) the subdivision benefit (Repp, 2003). The filled duration illusion refers to the tendency for the perceived duration of an inter-beat interval to increase when it is subdivided by additional metrical events, whereas the subdivision benefit refers to reduced SMS variability when beat subdivisions provide additional temporal reference points between successive beats (Repp, 2003; Repp and Dogget, 2007; Repp, 2010). The subdivision benefit has been proposed to arise from improved temporal prediction, potentially because additional temporal reference points reduce the perceptual underestimation of inter-onset intervals. Such perceptual underestimation may in turn explain why people tap slightly ahead of the beat during sensorimotor synchronization (Wohlschläger and Koch, 2000). A better understanding of the neural mechanisms underlying the subdivision benefit could help optimize rhythmic auditory stimulation for individuals with impaired motor coordination, including those with Parkinson’s disease (Damm et al., 2020).

However, although both musicians and non-musicians benefit from subdivisions at different ratios and tempi (Repp, 2008a, Repp and Bruttomesso, 2009), the magnitude of the subdivision benefit is not uniform across individuals but also depends on musical expertise. For example, Repp (2008a) replicated the subdivision benefit by showing that metrical subdivision reduced the variability of synchronization asynchronies. Extending these findings, Zendel et al. (2011) showed that improvements in both tapping variability and negative mean asynchrony were primarily explained by shorter inter-onset intervals (IOIs) introduced by subdivision rather than by longer inter-tapping interval (ITIs). However, musicians synchronized better than non-musicians with subdivisions ranging from 1:2 to 1:9 ratios, while non-musicians’ tapping synchronization was reduced for all uneven subdivisions (Repp, 2007). Tapping synchronization generally improves when participants imagine an antiphase beat (i.e., 1:2, the duplet) or tap to every other sound (i.e. 2:1, binary meter; see Repp and Dogget, 2007; Repp, 2010b), and latent expectations of binary subdivisions occur automatically when synchronizing to a beat (Repp, 2008b), but can flexibly be induced in a ternary subdivision context (Repp and Jendoubi, 2009). Furthermore, binary subdivisions are present in most cultures worldwide (Savage et al., 2015) and emerge spontaneously in iterated learning paradigms (Ravignani et al., 2016), likely due to categorical perception (Ravignani et al., 2018). Altogether, these studies suggest that binary groupings are pervasive in human cognition, and that musical training can improve SMS to non-binary subdivisions.

Beyond behavioral effects of beat subdivisions on tapping synchronization, previous research has begun to investigate the neural mechanisms underlying the perception of rhythmic structure using EEG frequency tagging (Nozaradan, 2014; Nozaradan et al., 2018). Frequency tagging measures neural activity associated with specific frequencies of an external stimulus by decomposing steady-state evoked potentials (SSEPs, i.e., neural responses that are frequency- and phase-locked to the stimulus) into amplitudes in the frequency domain. According to neural resonance theory (Large and Snyder, 2009), beat perception emerges through the synchronization of neural populations to rhythmic inputs, giving rise to oscillatory activity at the beat frequency as well as its harmonics and subharmonics that reflect subdivisions and higher-order metrical structure. Consistent with this framework, frequency-tagging studies have demonstrated neural responses that track beat-related periodicities and higher-order metrical groupings, providing evidence for internal neural representations of rhythm (Lenc et al., 2021; Harding et al., 2025).

Higher-order oscillations in EEG and MEG recordings further demonstrate how the brain integrates both bottom-up sensory input with top-down predictive processes during beat and meter perception (Fujioka et al., 2010; Nozaradan et al., 2011; Stupacher et al., 2016; Celma-Miralles et al., 2016). Importantly, neural synchronization to rhythmic stimulation (i.e., neural tracking, or more broadly termed as entrainment, Obleser and Kayser, 2019; see also Haegens, 2020; Doelling and Assaneo, 2021) is not merely a passive reflection of stimulus properties but also reflects endogenous neural processes that selectively enhance behaviorally relevant periodicities (Lenc et al., 2020). Such selective enhancement has been demonstrated for the beat of complex rhythms (Nozaradan et al., 2016; Tal et al., 2017), periodic pulses of naturalistic musical excerpts (Tierney and Kraus, 2015) and the hierarchical syntactic structure of linguistic phrases (Ding et al., 2017). Together, these findings suggest that selective enhancement of behaviorally relevant periodicities enables more efficient synchronization with rhythmic input (Gibbings et al., 2023) and supports accurate temporal predictions of the beat and metrical structure (Damsma et al., 2025).

Although prior studies established that subdivisions are fundamental to rhythm perception and synchronization, they have rarely been investigated directly as the experimental variable. Previous frequency-tagging studies have focused mainly on neural responses to slower metrical levels (Nozaradan, 2014; Vuust and Witek, 2014; Fujioka et al., 2015; Cirelli et al., 2016), which has limited the examination of subdivisions. Some studies have dissociated metrical structure from event periodicity by comparing binary and ternary groupings over the same auditory sequence (Bouvet et al., 2020) or by tapping to every other sound of the periodic sequence (Nozaradan et al., 2015). Although both metrical groupings and beat subdivisions are essential building blocks of rhythm perception and production, beat groupings occur on a slower time scale than subdivisions. Subdivisions are implicitly embedded in complex rhythmic structures, often generating binary and ternary groupings of events (Nozaradan et al., 2012), and building up syncopations (Tal et al., 2017) and polyrhythms (Vuust et al. 2006; Stupacher et al., 2017; Møller et al, 2026). These metrical organizations are reflected in both mismatch negativities (MMN; Vuust et al., 2005), that depend on metrically differentiated event-related potentials (ERP; Brochard et al., 2003; Bouwer et al., 2016, 2024), and frequency-tagged neural responses, that selectively enhance behaviorally relevant metrical periodicities (Celma-Miralles et al., 2024; Møller et al, 2026).

Despite converging behavioral and neurophysiological evidence for the importance of beat subdivisions, it remains unclear how the brain represents subdivisions as rhythmic events, rather than as implicit components of more complex rhythmic structures. Little is known about how subdivision ratio influences neural synchronization during active sensorimotor synchronization, how it shapes predictive processing of temporal events, and how these mechanisms are modified by musical expertise. To address these questions, we measured neural and behavioral responses while participants synchronized with the beat of four subdivisions. We combined frequency tagging with a finger-tapping paradigm to examine whether different subdivision ratios differentially modulate neural synchronization and sensorimotor synchronization in musicians and non-musicians. We hypothesized that listening to binary subdivisions (duplets and quadruplets) would produce more stable tapping performances than non-binary subdivisions (triplets and quintuplets), accompanied by stronger neural synchronization to the beat frequency and its harmonics, particularly in musicians due to their expertise in SMS. Finally, we analyzed ERPs to compare neural responses to unexpected sound onsets and their corresponding omissions, testing whether metrically less predictable subdivision timings elicit larger prediction errors (Bouwer et al., 2024).

## 2. MATERIALS and METHODS

### 2.1 Participants

Thirty-one healthy volunteers (22 female, mean age: 23.39 ±3.21, age range: 19-35) participated in this study after providing written informed consent. Sixteen of them pursued (at least) ten years of formal training in music and/or extensive musical practice (henceforth, *musicians*). The other fifteen participants pursued less than two years of basic music training, if any, (henceforth, *non-musicians*). No one of them reported any hearing, psychiatric or neurological disorder. Thus, the inclusion criteria depended on the years of music training of our healthy participants. The number of participants was selected based on the positive findings found in a preliminary EEG pilot with sixteen non-musicians (Farrokh, 2018). To validate the two groups, participants performed the rhythmic discrimination of the Musical Ear Test (MET, Wallentin et al., 2010), which revealed that the performance of musicians (M = 79.57, SE = 1.65) were better than the performance of non-musicians (M = 66.28, SE = 1.88; *t*_(29)_ = 5.33, *p* < .001). All volunteers were compensated with 100 Danish Kroner per hour for participating in the experiment.

The research was carried out in compliance with the ethical standards outlined in the Declaration of Helsinki and the Danish Code of Conduct for Research. Additionally, all procedures were approved by the Institutional Review Board of the Danish Neuroscience Center at Aarhus University (Denmark).

### 2.2 Auditory Stimuli

The stimuli consisted of isochronous sequences of tones, in which a periodic beat was subdivided into duplets (1:2), triplets (1:3), quadruplets (1:4) and quintuplets (1:5). The beat frequency was set at 1.25 Hz because previous research revealed that it easily elicits steady-state evoked potentials and allowed all subdivisions to fall within the ecological range of beat perception (London, 2012). We created a 30 ms-pure tone at the fundamental frequency of 659 Hz, whose amplitude was shaped from 0 to 1 with a sinusoidal bell curve (see Figure 1). To create each auditory rhythm, we presented this sound at distinct inter-onset intervals (IOI). There were five different rhythmic sequences that lasted 20 seconds, the beat (25 sounds presented at 1.25 Hz, IOI = 800 ms), the duplets (50 sounds at 2.5 Hz, IOI = 400 ms), the triplets (75 sounds presented at 3.75 Hz, IOI = 266 ms), the quadruplets (100 sounds presented at at 5 Hz, IOI = 200 ms), and the quintuplets (125 sounds presented at at 6.25 Hz, IOI = 160 ms). For the practice trials, we used shorter sound sequences that lasted 6.4 seconds, in which the sounds that did not occur at the beat frequency —those that created each subdivision rhythm and occupied offbeat positions— were reduced 50% in their amplitude. All the auditory stimuli were created in Matlab 2019b (MathWorks) and presented binaurally through in-ear headphones at a comfortable hearing level, by using Psychtoolbox.

**Figure 1.**
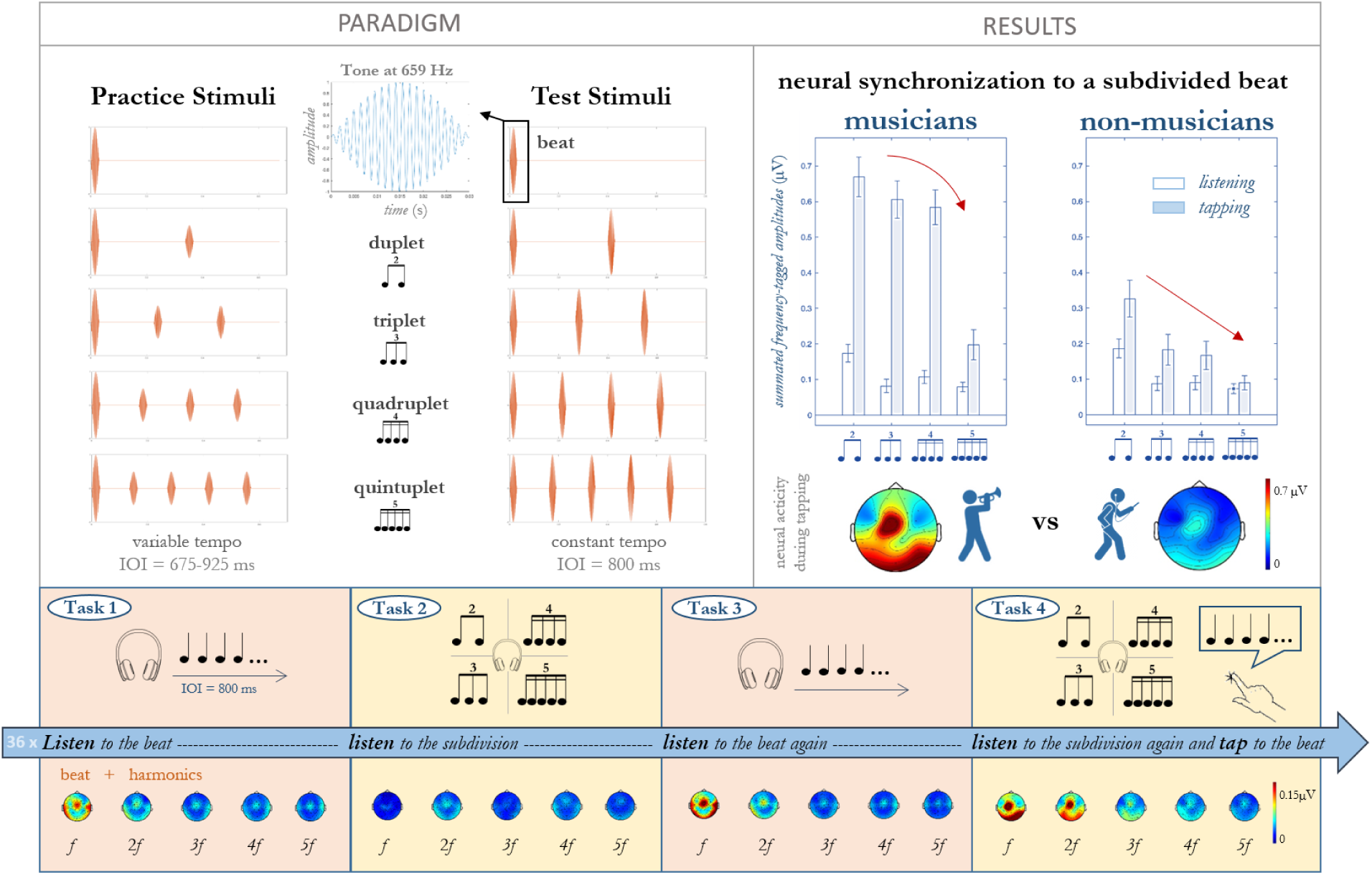
Our stimuli, paradigm and neural synchronization differences between musicians and non-musicians. The top left section depicts our 30-ms sound used to create the beat, the duplets, the triplets, the quadruplets and the quintuplets. A variable range of tempi and differences in amplitude were used for the training, while a constant tempo (IOI=800 ms, *f =* 1.25 Hz) and amplitude were used for the study. The bottom part depicts our paradigm, which involves four 20-second tasks for every auditory sequence: (1) listen to the beat, (2) listen to the subdivision, (3) listen to the beat, and (4) listen to the same subdivision while tapping to the beat. The neural synchronization of all participants is averaged and depicted for each task as topographies at each frequency of interest. The top right section shows musicians and non-musicians’ differences between listening and tapping to the subdivisions. Notice the greater increases in musicians for all subdivisions. This difference is also evident in the topographies below, which show the neural synchronization to beat-related frequencies averaging all subdivisions.

### 2.3 Paradigm

Participants were comfortably seated in a silent room at the Royal Academy of Music of Aarhus and completed two tasks. Firstly, they performed the rhythmic part of the Musical Ear Test (Wallentin et al., 2010), which involved listening to 52 pairs of rhythmic sequences and indicating on a paper form whether they were identical or different. During the EEG paradigm, participants faced a screen that displayed experimental instructions, alerts for two breaks (approximately 15 minutes each), and a fixation cross to keep their eyes still during the tasks. A tapping midi device (Arturia BeatStep) was positioned within easy reach in front of the participants. At the beginning of the task, participants were given short practice trials and instructed by the experimenter when to tap the beat of the distinct subdivisions. Each auditory trial had four parts: (1) listening to an isochronous beat, (2) listening to one of the four subdivisions, (3) listening to the beat again, and (4) listening to the same subdivision while tapping the beat (see Figure 1). Before the tapping task, the fixation cross changed into a synchronized count-down from 3 to 1, to warn and prepare participants. After completing eight practice trials, the experiment began, and participants listened to the actual trials presented in a pseudo-randomized order, repeated nine times. This order was counterbalanced across participants to prevent consecutive repetitions of the same trial. During the two breaks, the experimenter addressed any comments from participants and monitored the quality of the EEG recordings.

### 2.4 EEG recordings

We used the BrainVision Analyzer Software package (BrainVision Recorder 2.1, Gilching, Germany) with a BrainAmp amplifier (Brain Products GmbH, Germany) to record each participant’s electroencephalographic (EEG) signal. We placed a 32-electrodes actiCap (Fp1, Fp2, F3, F4, C3, C4, P3, P4, F7, F8, T7, T8, P7, P8, Fz, Cz, Pz, FC1, FC2, CP1, CP2, FC5, FC6, CP5, CP6, TP9, TP10, Oz) on the scalp according to the International 10/20 system. Two electrodes were placed on the left and right mastoid (RM and LM). Eye movements were tracked using two electrodes placed on the right eye: at the infra-orbital ridge and at the outer canthus (EOG-V and EOG-H). We recorded the signal using the FCz channel as a reference, keeping electrode impedances below 20 kΩ. The signal was amplified and digitized at a sampling rate of 1000 Hz, with an online low cut-off of 10 seconds and a high cut-off of 1000 Hz.

### 2.5 Behavioral analyses

Behavioral analyses were conducted to evaluate differences in finger-tapping performance between musicians and non-musicians. To this end, we computed the inter-tap intervals (ITIs) and tapping consistency. To correct for double-taps, we removed the second tap of two consecutive taps that occurred within ITIs less than 150 ms. We also adjusted the timings for an approximate recording delay of 20 ms. Additionally, we transformed the finger-tapping responses into a common angular scale in radians by multiplying each tap’s value by 2π and dividing by the beat IOI (800 ms). Using the Matlab Toolbox for Circular Statistics (Berens, 2009), we calculated the mean resultant vector and its length for each subject, trial, and subdivision rhythm. The angular direction of the mean resultant vector indicates the taps’ position (i.e., phase) relative to an isochronous beat, while the length of the mean resultant vector reflects how closely the taps cluster around the mean angular direction.

To verify that the taps were not randomly performed, an Omnibus test was conducted, which confirmed that the angular directions of all participants were not uniformly distributed around the circle (*p* < .001). Circular V-tests were then performed, which showed that the mean vector directions of both musicians and non-musicians pointed towards 0°, corresponding to the beat “onset,” for all subdivision rhythms (*p* < .001), except for quintuplets in non-musicians (*p* = .105).

To test for the effects of the subdivision rhythm and musical training on tapping accuracy, a robust circular two-factor ANOVA known as Harrison-Kanji test was performed (Berens 2007). To compare tapping accuracy across groups for each subdivision rhythm, Watson-Williams tests were used, which are circular statistical analogues of two-sample *t*-tests. To assess differences in tapping consistency, a standard Mixed-Design ANOVA was used on the mean vector length, with the within factor Rhythm (duplet, triplet, quadruplet, quintuplet) and the between-factor Group (musicians, non-musicians). This analysis was followed by Bonferroni-corrected post hoc *t*-tests. Additional non-parametric tests were reported in Supplementary Materials to account for ceiling values and confirm the results in each group.

### 2.6 EEG analyses

The EEG recordings were pre-processed using BrainVision Analyzer 2.1 (Brain Products GmbH). Any channel that appeared very noisy or flat was interpolated from the surrounding channels via spherical spline interpolation. Subsequently, all channels were re-referenced to a common average-reference, which yielded the channel FCz for subsequent analyses. To remove slow drifts in the recordings, a zero-phase Butterworth filter was applied with a notch filter at 50 Hz, a high pass filter at 0.1 Hz (with order 8) and a low pass filter at 30 Hz (with order 8). Offline, we automatically detected whether the EEG activity crossed a maximal allowed voltage step of 50 μV/ms, a maximal allowed difference of voltage of ±100 μV or a voltage lower than 0.5 μV, and marked it as bad intervals within ±200 ms. Eye blinks and clear muscular movements were removed from the signal by using the Ocular Correction ICA function, which was based on the EEG data free of bad intervals. Finally, the filtered EEG data were segmented based on task and subdivision rhythm, and the nine 20-second trials were exported into European Data Format. All further analyses were performed using Matlab (The MathWorks, version 2023b), JASP software (JASP Team 2025, version 0.95.4.), and R Studio software (version R-3.6.2).

To eliminate evoked potentials related to stimuli onset and changes across tasks, the first 1.6 seconds of each 20-second sequence were removed, as steady-state evoked potentials require some repetitions to appear (Nozaradan et al., 2011). EEG epochs were then averaged for each participant and task across nine trials to reduce non-phase-locked neural activity to the stimuli or to the motor task. The EEG signal underwent fast Fourier transform (zero-padded to the next power of two: 2^15) and the amplitude (in μV) was obtained over a frequency range from 0 to 500 Hz and a bin frequency resolution of 0.0305 Hz. The amplitudes were the sum of neural activity induced by processing the stimuli and some background noise or spontaneous activity. To increase the signal-to-noise ratio, the averaged amplitude of the two surrounding non-adjacent frequency bins (-0.153 to -0.122 Hz and 0.122 to 0.153 Hz) was subtracted from each frequency bin. If no steady-state evoked potentials were present, all frequencies of the spectra should vary similarly (Nozaradan, 2014). Therefore, after the subtraction, each frequency peak should randomly tend towards zero. The presence of peaks related to beat and its subdivisions in the frequency spectra was discovered by averaging all participants’ magnitudes centered at our frequencies of interest across the scalp and subjecting them to *t*-tests against zero (Table S1). Figure 2 shows these peaks on the frequency dimension, while Figure S1 depicts the values as scatter box violin plots. Table S1 reports the *t-*values and *p*-values of the t-tests for each frequency and subdivision. In Figure S2 we depict similar frequency spectra but obtaining the neighbor-subtracted amplitudes after applying fast Fourier transform to each single trial before averaging them (see Møller et al., 2026). This analysis might be noisier because non-phase locked activity is not reduced by averaging trials over time, but it has the potential to reveal neural synchronization to the beat when this is anchored at other sounds of the subdivisions (e.g., the second sound of the duplet).

**Figure 2.**
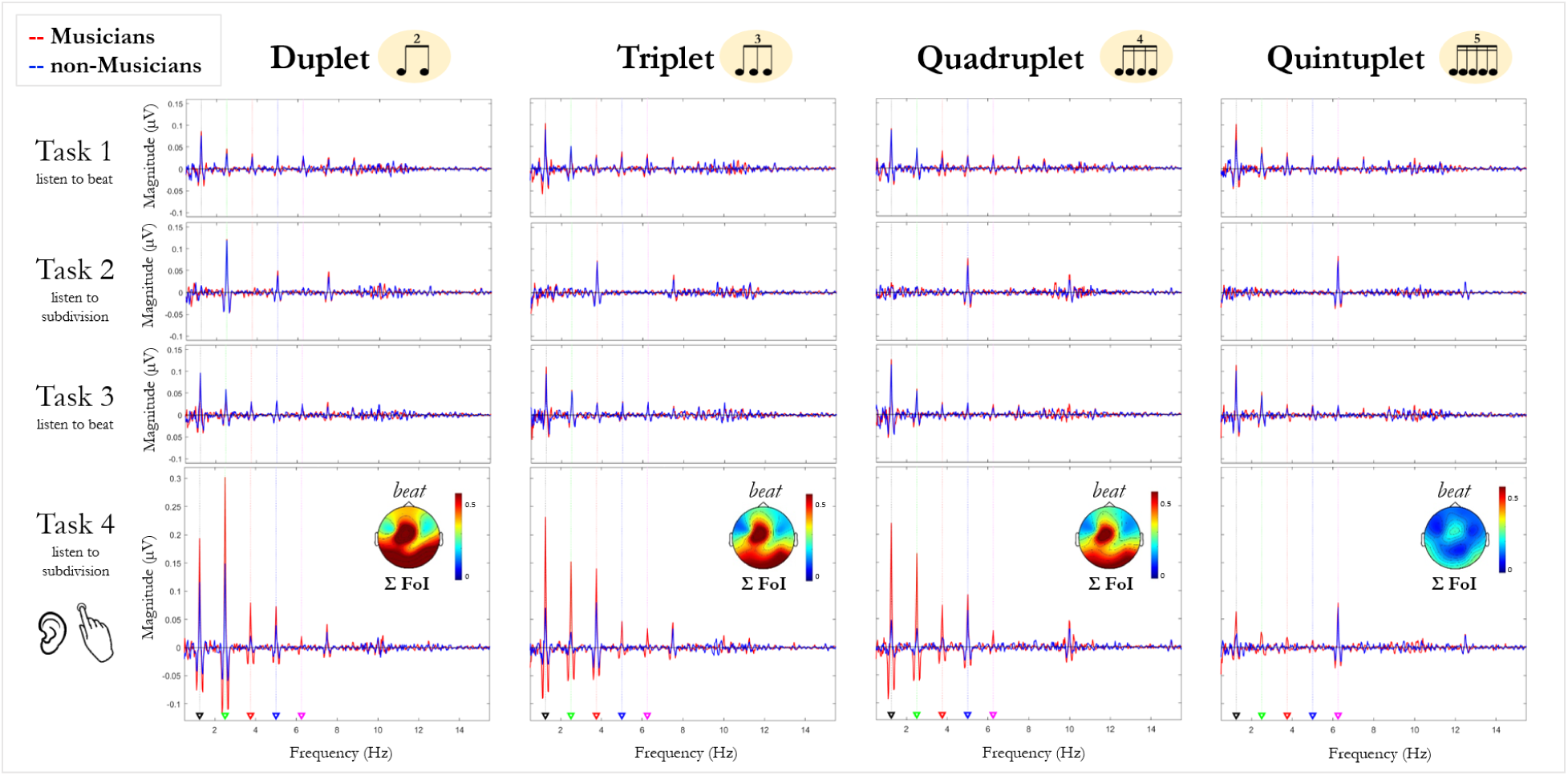
Frequency amplitudes of musicians and non-musicians for each task and subdivision. The red line stands for musicians’ mean, while the blue line stands for non-musicians mean at each task (rows) and subdivision (columns). The triangles signal the frequencies of the beat and its harmonics: black for the beat (*f*), green for the duplet (2*f*), red for the triplet (3*f*), blue for the quadruplet (4*f*), pink for the quintuplet (5*f*). At task 4, tapping to the beat while listening to the subdivision, topographies for all participants show the summated amplitudes at the five frequencies of interest. Frontocentral activity is slightly left-lateralized, as the participants performed the tapping task with their contralateral right hand.

The EEG analysis aimed to answer whether the four subdivisions elicit different neural activity in each task. To answer this question, we followed Retter et al. (2021) and summated the frequency-tagged amplitudes occurring at the five frequencies of interest (f, 2f, 3f, 4f and 5f, depicted with distinct colors in Figure 4; see also Figure S1). This should account for all neural activity related to the beat and the subdivisions, which correspond to the first 4 harmonics of the beat. For each task, a two-way Mixed-Design ANOVAs was applied to the summated peak amplitudes, with the within factor Rhythm (Duplets, Triplets, Quadruplets, Quintuplets) and the between factor Group (Musicians, non-Musicians). We corrected the degrees of freedom using Greenhouse-Geisser estimates of sphericity when the assumptions for sphericity were violated. When significant effects and/or interactions appeared, we conducted post hoc two-tailed *t*-tests with the Bonferroni correction (with significance level at *p* < .05) for all analyses.

To assess the impact of subdivisions on perceiving the isochronous beat (“listening to the beat” tasks), we subtracted Task 1 amplitudes from Task 3 amplitudes for each participant and subdivision. Similarly, to evaluate the contribution of tapping to the beat on listening to subdivisions, we subtracted Task 2 amplitudes from Task 4 amplitudes for each participant and subdivision. We performed one-sample *t*-tests to compare the subtracted neural amplitudes to zero. If data was not normally distributed according to the Shapiro-Wilk test of normality, we used Wilcoxon signed-rank tests instead of Student *t*-tests.

To examine the association between SMS and beat-related neural activity, we calculated the correlation between tapping consistency values and the summated frequency-tagged amplitudes of the subdivisions for each group. When data violated the assumption of normality (Shapiro-Wilk tests), Spearman’s rho was used instead of Pearson’s r.

To investigate ERP differences related to predictive processes and subdivision expectancies, we computed the difference waveforms to measure mismatch responses (Näätänen et al., 2007; Bouwer et al., 2024) related to the presentation or omission of first sound subdividing the beat differently during the listening tasks. Obligatory auditory evoked potentials (e.g. N1 responses) should be smaller for metrically expected than unexpected sound onsets to silence predicted sensory inputs (Lange, 2013), while expectancy violations (e.g., the attended MMN homologue N2b and the P300) should be larger for unexpected events that interfere with the perceived beat (Bouwer et al., 2024). First, we calculated ERPs for each beat cycle by segmenting all the trials into 800 ms. Secondly, we averaged frontocentral electrodes (i.e. Fz, FCz, Cz, FC1 and FC2) after centering their amplitudes around zero using an inside epoch baseline from 0 to 100 ms, since it homogeneously includes the first beat onset across tasks. For each task and subdivision, we then averaged beat cycles to calculate two ERP waveforms (see Fig. S3): the first beat cycle, referred to hereafter as the deviant epoch because unexpected sounds or omissions occurred only during this cycle, and the average of the remaining beat cycles, referred to as the standard epoch because they reflected the repeated stimulus pattern after habituation. Bad trials (i.e., voltage range exceeding 100 µV or voltage steps greater than 50 µV) were removed. For each subdivision, difference waves were calculated by subtracting the standard epoch of Task 2 from the deviant epoch of Task 2 or Task 3 to isolate N2b and P300 responses. Negative deflections followed by positive deflections appeared in the averaged ERPs (see Figure S3) and difference waves (Figure 6). Negative peak and positive peak latencies were respectively identified within the range 60-180 and 180-300 ms. Following Bouwer et al., 2024), mean amplitudes were calculated averaging the voltage over a 60-ms time window centered at the global peak latency of each subdivision. One sample *t*-tests against zero verified the presence of N2b and P300 responses (see Table S2), and Mixed-Design ANOVAs compared them across Subdivisions and Groups. Greenhouse-Geisser corrections were applied when sphericity assumptions were violated, and Bonferroni adjustments corrected significance levels for multiple comparisons in post hoc *t*-tests.

## 3. RESULTS

### 3.1 Finger tapping accuracy

A Harrison-Kanji test revealed an effect of Group (F_(1, 123)_ = 13.11, *p* < .001) and an effect of Rhythm (F_(3, 123)_ = 3.29, *p* = .023), but no interaction between them (F_(3, 123)_ = 1.65, *p* = .181). The post-hoc Watson-Williams tests revealed that there were differences between musicians and non-musicians (F_(1, 123)_ = 5.55, *p* < .020), but no differences between the four subdivision rhythms when all participants are averaged (all *p* >.05).

Although the Group × Rhythm interaction was not significant, exploratory Watson– Williams tests were conducted to characterize potential differences between musicians and non-musicians within individual subdivision rhythms. These comparisons revealed a group difference for quintuplets (*p* = .005). Furthermore, the angular directions during tapping to quintuplets did not differ from the other subdivision rhythms in musicians (all *p* > .05), whereas in non-musicians they differed from triplets (*p* = .025) and quadruplets (*p* = .038), but not from duplets (*p* = .073). We do not include these accuracy results in the Discussion.

Overall, musicians showed more accurate synchronization than non-musicians, whereas subdivision rhythm had only a limited influence on tapping accuracy. Exploratory analyses suggested that group differences were most pronounced for quintuplets (Figure 3a).

**Figure 3.**
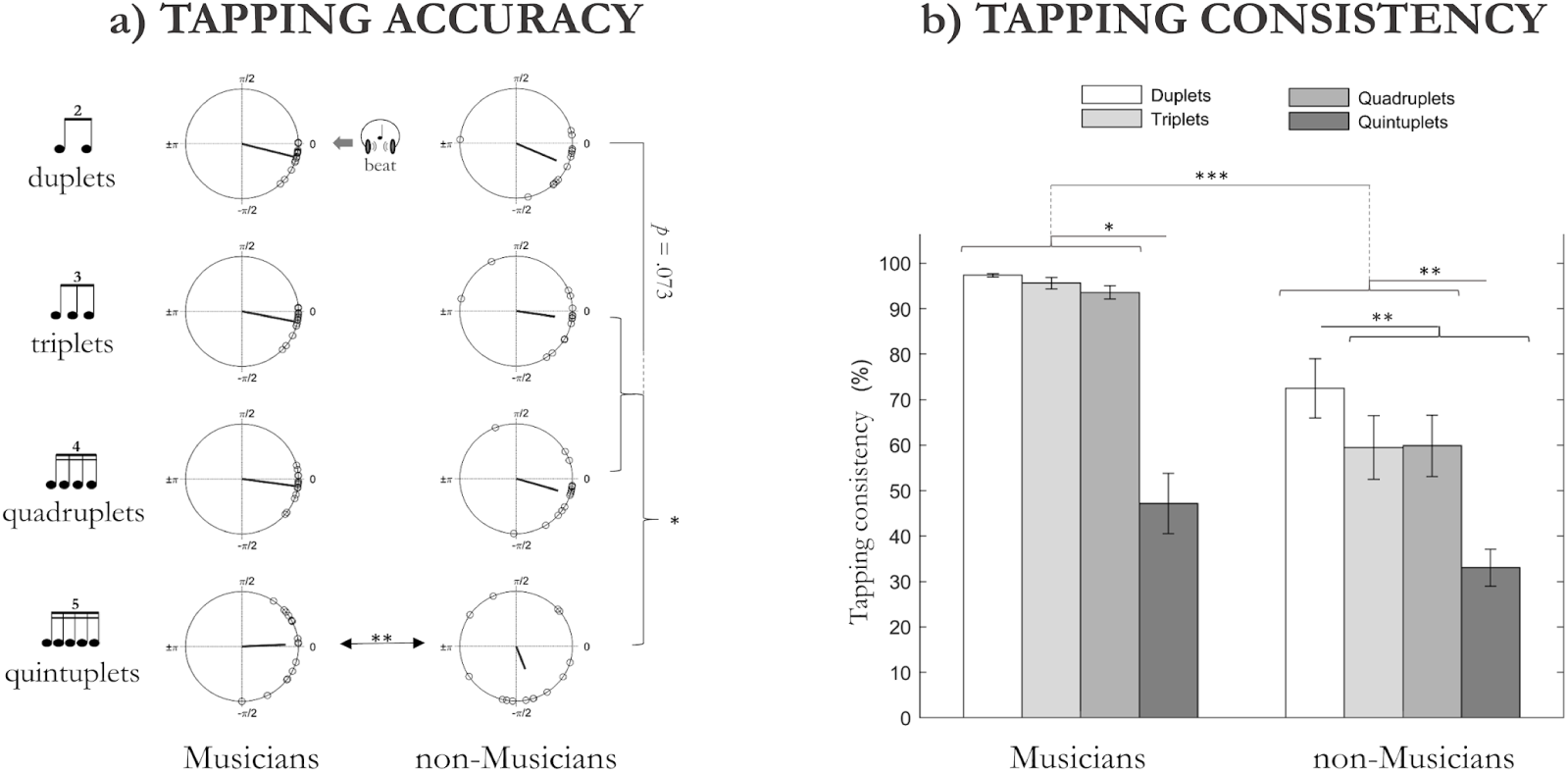
Tapping responses for each group and subdivision. The rose-plots (a) represent non-musicians’ and musicians’ tapping accuracy for duplets, triplets, quadruplets and quintuplets. The downbeat of the subdivision should be located close to 0°. The bar graph (b) represents non-musicians’ and musicians’ tapping consistency for duplets, triplets, quadruplets and quintuplets. * stands for *p* < .05; ** stands for *p* < .01; *** stands for *p* < .001.

### 3.2 Finger tapping consistency

A Mixed-Design Anova revealed a main effect of Rhythm (F_(1.63, 47.28)_ = 69.05, *p* < .001), a main effect of Group (F_(1, 29)_ = 23.80, *p* < .001) and an interaction between them (F_(1.63, 47.28)_ = 4.22, *p* = .027). Bonferroni-corrected post hoc *t*-tests (see Figure 3b) revealed that musicians exhibited higher tapping consistency than non-musicians for all rhythmic subdivisions (all *p* < .001) except quintuplets (*p* = .083). Within-group comparisons further revealed distinct patterns. For non-musicians, duplets (M = 0.725, SE = 0.253) had higher consistency than triplets (M = 0.595, SD = 0.272), quadruplets (M = 0.599, SD = 0.262) and quintuplets (M = 0.330, SD = 0.158, all *p* ≤ .005), whereas quintuplets had lower consistency than duplets, triplets and quadruplets (all *p* ≤ .002). For musicians, only quintuplets (M = 0.472, SD = 0.264) had lower consistency than duplets (M = 0.974, SD = 0.014), triplets (M = 0.957, SE = 0.050) and quadruplets (M = 0.936, SD = 0.060, all *p* < .001). Since the tapping consistency of musicians for duplets, triplets and quadruplets reached almost ceiling values that skewed the normal distribution, we also applied a non-parametric ANOVA to each group to confirm that ceiling effects did not obscure subdivision-related differences. Friedman tests and Conover’s post hoc comparisons confirmed the parametric results across subdivisions (see Tables S7 and S8).

In summary, tapping consistency was higher in musicians than non-musicians across rhythmic subdivisions. Whereas musicians maintained consistently high performance for duplets, triplets, and quadruplets, non-musicians showed a selective advantage for duplets and a marked reduction in consistency for quintuplets (Figure 3b).

### 3.3 Steady-state Evoked Potentials related to the beat and its subdivisions

For the two “listening to the beat” tasks there were no significant main effects of Rhythm or Group (all *p* > .05) in the 2-way Mixed-Design ANOVAs. For the “listening to the subdivision” task, there was a significant main effect of subdivision (F_(3,87)_ = 17.027, *p* < .001, η²_p_ = 0.370). Further post hoc *t-*tests revealed that duplets elicited higher neural activity (M = 0.180, SD = 0.100) than triplets (M = 0.085, SD = 0.075, *p* < .001), quadruplets (M = 0.099, SD = 0.074, *p* < .001) and quintuplets (M = 0.076, SD = 0.053, *p* < .001). For the “listening to the subdivision and tapping to the beat” task, there were significant main effects of Rhythm (F_(1.99,57.81)_ = 46.089, *p* < .001, η²_p_ = 0.614) and Group (F_(1,29)_ = 37.671, *p* < .001, η²_p_ = 0.565) and the interaction Rhythm*Group (F_(1.99,57.81)_ = 11.277, *p* < .001, η²_p_ = 0.280). Musicians had lower neural amplitudes for quintuplets (M = 0.198, SD = 0.169) , than duplets (M = 0.670, SD = 0.224, *p* < .001), triplets (M = 0.606, SD = 0.211, *p* < .001), and quadruplets (M = 0.584, SD = 0.194, *p* < .001), while non-musicians had higher amplitudes for duplets (M = 0.326, SD = 0.200), than triplets (M = 0.183, SD = 0.168, *p* < .001), quadruplets (M = 0.167, SD = 0.153, *p* < .001) and quintuplets (M = 0.090, SD = 0.078, *p* < .001). Musicians had higher amplitudes than non-musicians for all subdivision rhythms (all *p* < .001) except for quintuplets (*p* > .050). See Figure 4b for a graphical representation of this interaction.

**Figure 4.**
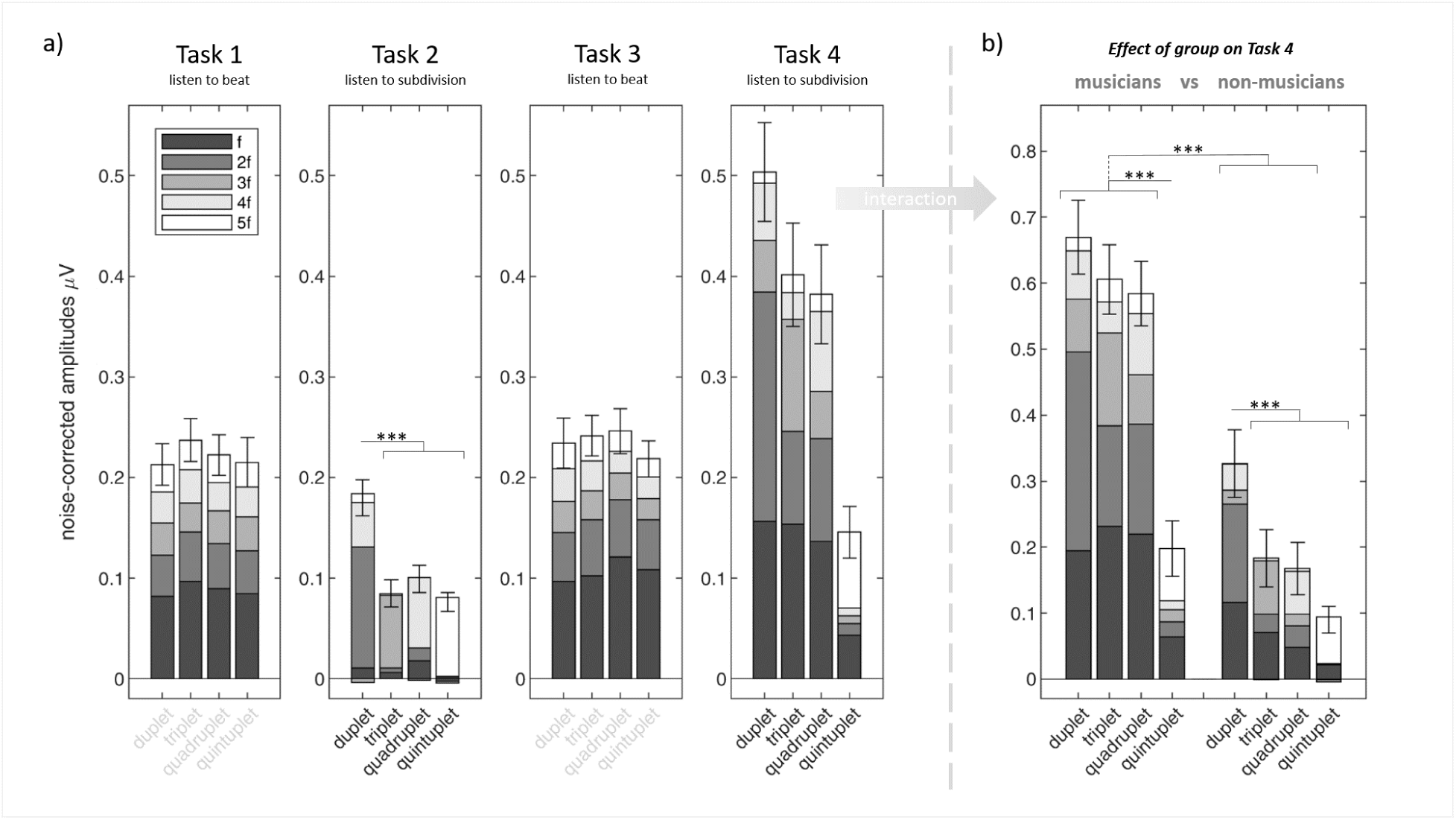
Frequency-tagged responses to each task of the sequence including a distinct subdivision: duplets, triplets, quadruplets and quintuplets. The bar graphs with SEM confidence intervals depict the summation of noise-corrected frequency-tagged amplitudes at the five frequencies of interest: the beat (f) and its harmonics (2f, 3f, 4f, 5f) color-coded in grayscale. (a) The effect of subdivision in the four listening tasks of the auditory sequence comparing the neural amplitudes. (b) The interaction between group and subdivision in the “listening to subdivision task with sensorimotor synchronization”, in which musicians and non-musicians tapped to their underlying beat. Listening to duplets and finger-tapping to their underlying beat seem to elicit greater neural synchronized activity across groups. Notice that, when listening to subdivision, duplets are dominated by 2f, triplets by 3f, quadruplets by 4f and quintuplets by 5f. Since in task 1 and Task 3 participants listened to the same beat, we dimmed the labels of the subdivisions. Statistical significance is denoted by *** (*p* < .001).

In summary, duplets were associated with the largest summed frequency-tagged amplitudes during passive listening across both groups and during synchronized tapping in non-musicians. In contrast, musicians showed reduced summed amplitudes for quintuplets during synchronized tapping, resulting in no significant group differences for this subdivision.

To identify the contribution of individual frequency components underlying the summed frequency-tagged amplitudes, Supplementary Material (Section 4) reports analyses in which each frequency of interest was examined separately. These analyses revealed increased amplitudes at the beat frequency and its first harmonic across all subdivision rhythms in both musicians and non-musicians (Figures S4 and S6a).

### 3.4 Comparisons of the neural amplitudes between tasks for each subdivision and group

One sample *t*-tests comparing the difference in summed frequency-tagged amplitudes between the two listening to the beat tasks (Task 3 − Task 1) revealed no significant results (all *p* > .050). In contrast, comparisons between the listening to the subdivision task and the listening to the subdivision while tapping to the beat task (Task 4 − Task 2) revealed significant increases in summed amplitudes (Table 1). Musicians showed significant increases across all subdivision rhythms relative to passive listening (all *p* < .001). In non-musicians, summed amplitudes also increased during synchronized tapping, but only duplets and triplets reached uncorrected significance, and neither survived Bonferroni correction.

**Table 1.** One sample *t*-tests against zero for the neural amplitudes resulting from the subtraction Task 4 - Task 2, for each subdivision and group. Wilcoxon signed-rank tests are reported, instead of Student *t*-tests, when ShapiroWilk tests of Normality indicate that the data is not normally distributed. Reported *p*-values are not corrected for multiple comparisons: a strict Bonferroni correction would reject the results found for duplets and triplets in non-musicians.

| Task4-Task2 | duplets |  | triplets |  | quadruplets |  | quintuplets |  |
| --- | --- | --- | --- | --- | --- | --- | --- | --- |
| musician<br>s | Student *t* | *p*-value | Student *t* | *p*-value | Student *t* | *p*-value | Wilcoxon | *p*-value |
|  | 10.249 | <.001 | 9.712 | <.001 | 9.698 | <.001 | 129 | <.001 |
| non-musician<br>s | Wilcoxon | *p*-value | Wilcoxon | *p*-value | Student *t* | *p*-value | Student *t* | *p*-value |
|  | 104 | .010 | 99 | .026 | 2.129 | .052 | 1.184 | .256 |

### 3.5 Correlations between tapping and SS-EPs

Table 2 summarizes the correlations between tapping consistency and frequency-tagged amplitudes during synchronized tapping. Significant positive correlations were observed when all participants were analyzed together, although separate analyses revealed distinct patterns across groups. In musicians, tapping consistency correlated positively with summed amplitudes only for quintuplets. In contrast, non-musicians showed positive correlations for duplets, triplets, and quadruplets. However, after, the correlation for duplets in non-musicians and the correlation for quintuplets in the pooled sample do not survive Bonferroni correction for multiple comparisons. no longer significant

**Table 2.** Correlations between the consistency of the tapping responses and the neural amplitudes during tapping for each group and subdivision. Spearman’s rho is reported in not normally-distributed pairs. Reported *p*-values are not corrected for multiple comparisons: a strict Bonferroni correction would reject the correlation found for duplets in non-musicians and quintuplets in “all participants”.

|  | duplets |  | triplets |  | quadruplets |  | quintuplets |  |
| --- | --- | --- | --- | --- | --- | --- | --- | --- |
| “all participants” | Spearman's rho | *p-value* | Spearman's rho | *p-value* | Spearman's rho | *p-value* | Spearman's rho | *p-value* |
|  | 0.611 | <.001 | 0.812 | <.001 | 0.768 | <.001 | 0.452 | .011 |
| musicians | Spearman's rho | *p-value* | Spearman's rho | *p-value* | Pearson's r | *p-value* | Spearman's rho | *p-value* |
|  | 0.212 | .430 | 0.256 | .338 | 0.247 | .356 | 0.791 | <.001 |
| non-musicians | Spearman's rho | *p-value* | Spearman's rho | *p-value* | Pearson's r | *p-value* | Pearson's r | *p-value* |
|  | 0.604 | .020 | 0.814 | <.001 | 0.790 | <.001 | 0.046 | .870 |

### 3.6 ERP comparisons from difference waveforms: N2b and P300

Regarding the difference wave for Task 2 (unexpected sound onsets), one-sample *t*-tests confirmed the presence of N2b. The ANOVA revealed a significant main effect of Subdivision (F_(2.51, 72.91)_ = 7.65, *p* < .001) but no main effect of Group (F_(1, 29)_ = 1.65, *p* = .209) or Subdivision × Group interaction (F_(2.51, 72.91)_ = 1.26, p = .294). Post hoc comparisons showed that duplets elicited significantly smaller N2b amplitudes than triplets (*t_(29)_ = 3.423, p* = .011), quadruplets (*t_(29)_ = 4.402, p* < .001), and quintuplets (*t_(29)_ =3.221, p* = .019), whereas no other pairwise comparisons became significant (all *p* > .050). One-sample *t*-tests also confirmed the presence of P300. However, the ANOVA revealed no significant main effect of Subdivision (F_(2.22, 64.28)_ = 0.36, *p* = .720), no significant Subdivision × Group interaction (F_(2.22, 64.28)_ = 0.98, *p* = .387), and no significant main effect of Group (F_(1, 29)_ = 3.37, p = .077). Although this effect did not reach significance, P300 amplitudes tended to be larger in musicians than in non-musicians (Mean Difference = 1.528, *t_(29)_* = 1.835, *p* = .077). Similar ANOVAs applied to individual peak latencies revealed no significant differences across subdivisions or groups (all *p* > .050), although there was a trend toward earlier N2b peaks in musicians (Mean Difference = 9.986, *t_(29)_* = 1.920, *p* = .065).

Regarding the difference wave for Task 3 (unexpected sound omissions), one-sample t-tests did not provide evidence for the presence of N2b. Consistent with this finding, the ANOVA revealed no significant main effects of Subdivision (F_(3, 87)_ = 0.81, *p* = .492) or Group, (F_(1, 29)_ = 0.35, *p* = .557), and no significant Subdivision × Group interaction (F_(3, 87)_ = 0.17, p = .920). In contrast, one-sample *t*-tests confirmed the presence of P300 (Table S2). The ANOVA revealed a significant main effect of Subdivision (F_(3, 87)_ = 3.14, *p* = .029), a trend in the main effect of Group, F(1, 29) = 3.11, *p* = .088, a significant Subdivision × Group interaction (F_(3, 87)_ = 2.998, *p* = .035). None of the between-group pairwise comparisons within individual subdivisions survived correction for multiple comparisons (all *p* > .050). Nevertheless, there was a trend toward larger P300 amplitudes for quintuplets in musicians than in non-musicians (Mean Difference = 3.419, *t*_(29)_= 3.15, *p* = .061; Figure 6b). Within musicians, P300 amplitudes were larger for quintuplets than for triplets (Mean Difference = 2.826, *t*_(29)_= 3.52, *p* = .023). Similar ANOVAs applied to individual peak latencies revealed no significant differences in N2b or P300 latencies across subdivisions or groups (all *p* >.050).

In sum, responses to unexpected sound onsets (i.e., Task 2, listening to the first subdividing event) primarily differed at the N2b level, where the significant effect revealed that duplets elicited the smallest responses across participants, while responses to unexpected sound omissions (i.e., Task 3, missing the first subdividing event) were mainly reflected in P300 amplitudes, where the significant interaction revealed the strongest responses for quintuplets in musicians (Figure 6).

## 4. DISCUSSION

Frequency tagging analyses revealed stronger neural synchronization to the beat and its harmonics in musicians than in non-musicians during synchronized tapping, whereas no group differences were observed during passive listening. Behaviorally, non-musicians showed a tapping advantage for the duplet subdivision relative to the other subdivisions, consistent with previous studies on subdivision benefits (Zendel et al. , 2011; Repp, 2003; Repp, 2007) and tapping preferences for binarized beats in polyrhythms (Møller et al., 2021; Stupacher et al., 2026; Møller et al., 2026). Listening to duplets increased beat-related amplitudes in both groups. In addition, ERP analyses showed that the N2b elicited by the first subdivision onset was smaller for duplets than for the other subdivisions, suggesting that binary subdivision was more consistent with listeners’ metrical expectations. Finally, musicians showed higher tapping consistency and stronger frequency-tagged responses than non-musicians for all subdivisions except quintuplets. This finding suggests that long-term musical training improves sensorimotor synchronization for the subdivisions most prevalent in Western music. Consequently, extensive experience with triplets and quadruplets may have diminished the behavioral and neural advantage for duplets that was observed in non-musicians.

### 4.1 Neural activity during tapping

Our study shows that sensorimotor synchronization to a subdivided beat increases neural synchronization to the beat and its harmonics, depending on musical training and the number of subdivisions, closely paralleling the behavioral results. Specifically, musicians showed increased neural amplitudes when tapping to the beat of the subdivisions (Figure 5b), consistent with their greater ability to synchronize movements metrically grouping the differently subdivided beat to keep the same pace. However, their amplitudes decreased for quintuplets (in line with Repp, 2007), indicating a potential limitation in their ability to synchronize to these faster and less common rhythms. In contrast, non-musicians exhibited subtle increases in neural amplitudes only for duplets and triplets. These are the simplest integer subdivision ratios found in music across cultures (Savage et al., 2015), commonly reproduced by both adults and children without formal musical training (Drake, 1998). Thus, non-musicians seemed to benefit primarily from binary and ternary subdivisions, while musicians also benefited from quadruplets. Both groups, however, showed lower amplitudes and worse tapping to the beat of quintuplets, indicating a general disadvantage for SMS to this subdivision, possibly due to its faster presentation rate and relative rarity in Western music.

**Figure 5.**
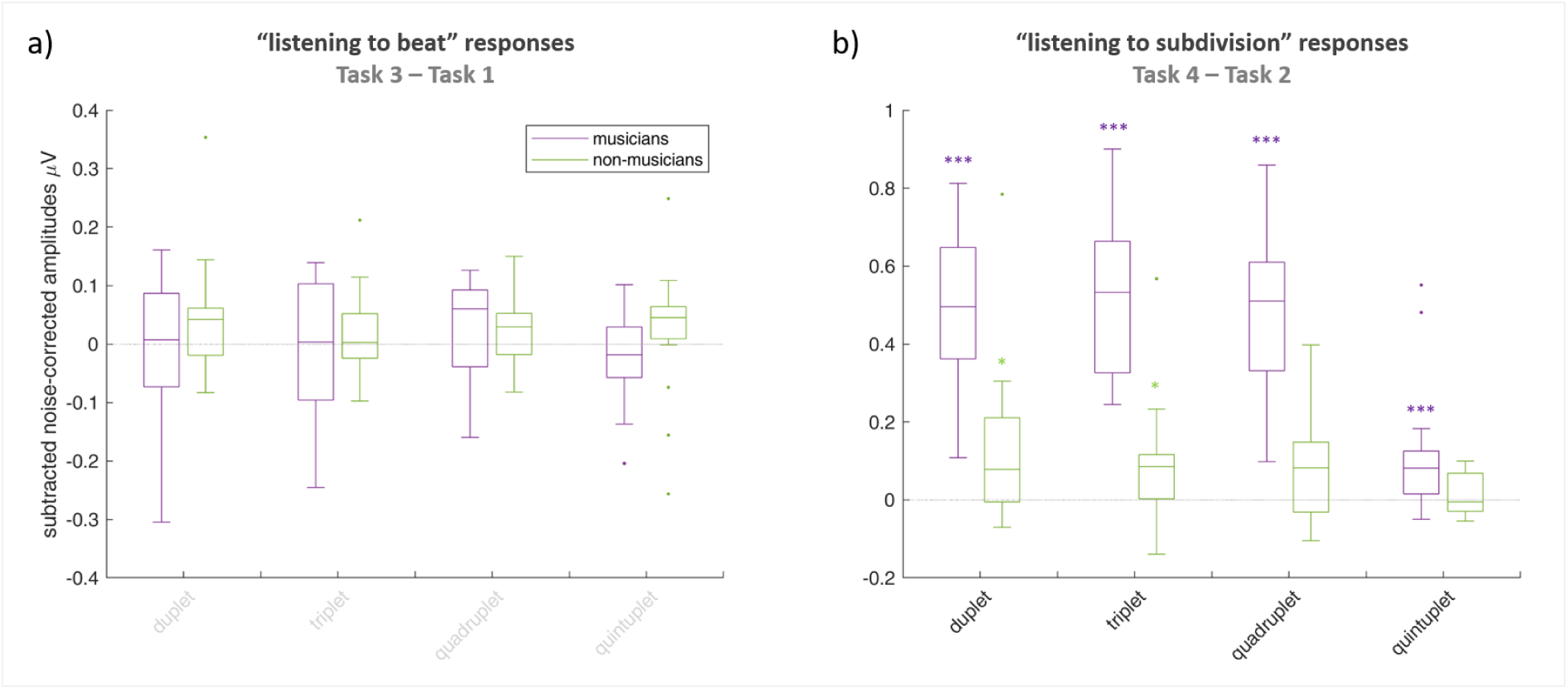
Difference in the frequency-tagged amplitudes between tasks with the same auditory stimuli. For each group, the boxplots depict the subtracted noise-corrected frequency-tagged amplitudes at each subdivision. (a) The neural amplitudes for Task 3 (“listening to beat” after presenting the subdivision) after subtracting Task 1 (“listening to beat” before presenting the subdivision). No comparison differed from zero. (b) The neural amplitudes for Task 4 (“listening to subdivision with sensorimotor synchronization”) after subtracting Task 2 (“listening to subdivision”), reflecting the contribution of tapping the underlying beat. While the amplitudes for musicians (in purple) increased for all subdivisions during tapping, the amplitudes for non-musicians (in green) only increased for duplets and triplets. Since in task 1 and Task 3 participants listened to the same beat, we dimmed the labels of the subdivisions. Outliers are depicted as a dot. Statistical significance is denoted by * for *p* < .05 and *** for *p* < .001.

Interestingly, neural responses differed between musicians and non-musicians only during active sensorimotor synchronization, and these differences depended on the subdivision rhythm. As mentioned above, the parallel changes in tapping consistency and neural amplitudes raise the possibility that increased neural synchronization partly reflects the consistency of the tapping responses and its corresponding somatosensory feedback. The reported correlations between neural amplitudes and tapping consistency support this idea when all participants are analyzed together, although this relationship appears to be primarily driven by the non-musician group. In contrast, the absence of significant correlations in musicians suggests that their frequency-tagged responses cannot be explained solely by synchronized motor activity. Instead, they may reflect stronger temporal predictions together with the integration of auditory and somatosensory information. Previous work likewise reported differences in neural synchronization between musicians and non-musicians in tasks that required top-down processing of the beat, such as maintaining a mental pulse (Stupacher et al., 2017), building a metrical structure (Celma-Miralles and Toro, 2019), or performing SMS tasks (Nozaradan et al., 2015; Lenc et al., 2020). Surprisingly, passive listening to the subdivisions did not produce group differences, despite the expectation that musicians would show stronger neural tracking of the beat underlying duplets, triplets, and quadruplets before performing the tapping task.

### 4.2 Neural activity when listening to duplets

The results from frequency tagging showed an unexpected increase in the summed frequency-tagged amplitudes during the passive *listening to subdivision* tasks when duplets were presented. One explanation is that binary subdivisions (1:2 ratio) are intrinsically easier to process and align with the underlying beat. This interpretation would be consistent with the small-integer ratio bias in timing (Ravignani et al., 2018; Mathias et al., 2020) and auditory perception (di Stefano et al. 2026), the widespread occurrence of binary structures across musical cultures (Savage et al., 2015), their spontaneous emergence in repeated melodic patterns (Celma-Miralles et al., 2024) and their proposed role as perceptual attractors for rhythm categorization (Barbero et al., 2024). However, an alternative explanation could be methodological: the larger amplitudes observed for duplets could reflect tempo-dependent modulations of the steady-state evoked potentials underlying beat perception (Damsma et al., 2025) or simply result from including an additional beat harmonic (4f) in the summed frequency-tagged amplitudes. To address this possibility, we analyzed each harmonic separately (Supplementary Materials, Section 4). These analyses showed no group differences across subdivisions during passive listening (Figure S5). Instead, they revealed increased amplitudes at the beat frequency and its first harmonic after listening to all subdivision rhythms (Figures S4 and S6). This pattern could reflect a tendency for the neural representation of rhythmic patterns to converge toward prototypical 1:1 and 1:2 ratios (Barbero et al., 2024), thereby simplifying and stabilizing beat perception following the subdivision period, and/or a general attentional gain to prepare for the sensorimotor synchronization period (De Pretto et al., 2018; Vogt 2009; Leech and Sharp 2014; Tierney and Kraus, 2014).

The ERP analyses provided converging evidence for a preferential processing of binary subdivisions, as reflected in neural responses associated with expectancy violations (Fig 6). Specifically, the N2b elicited by the first unexpected subdivision onset was smaller for duplets than for all other subdivision rhythms. This finding suggests that binary subdivision imposed lower processing demands because it was more consistent with listeners’ metrical expectations (Lange, 2003; Bouwer et al., 2024), in agreement with predictive coding of music (Vuust et al., 2022). In contrast, more complex subdivisions elicited greater mismatch responses. Although the evidence was not statistically conclusive, several ERP measures also pointed toward an effect of musical expertise. N2b peak latencies tended to occur earlier in musicians, while P300 amplitudes tended to be larger, both consistent with more precise temporal predictions in the musically trained brain. However, given the limited statistical power of the present study, these findings should be interpreted cautiously.

**Figure 6.**
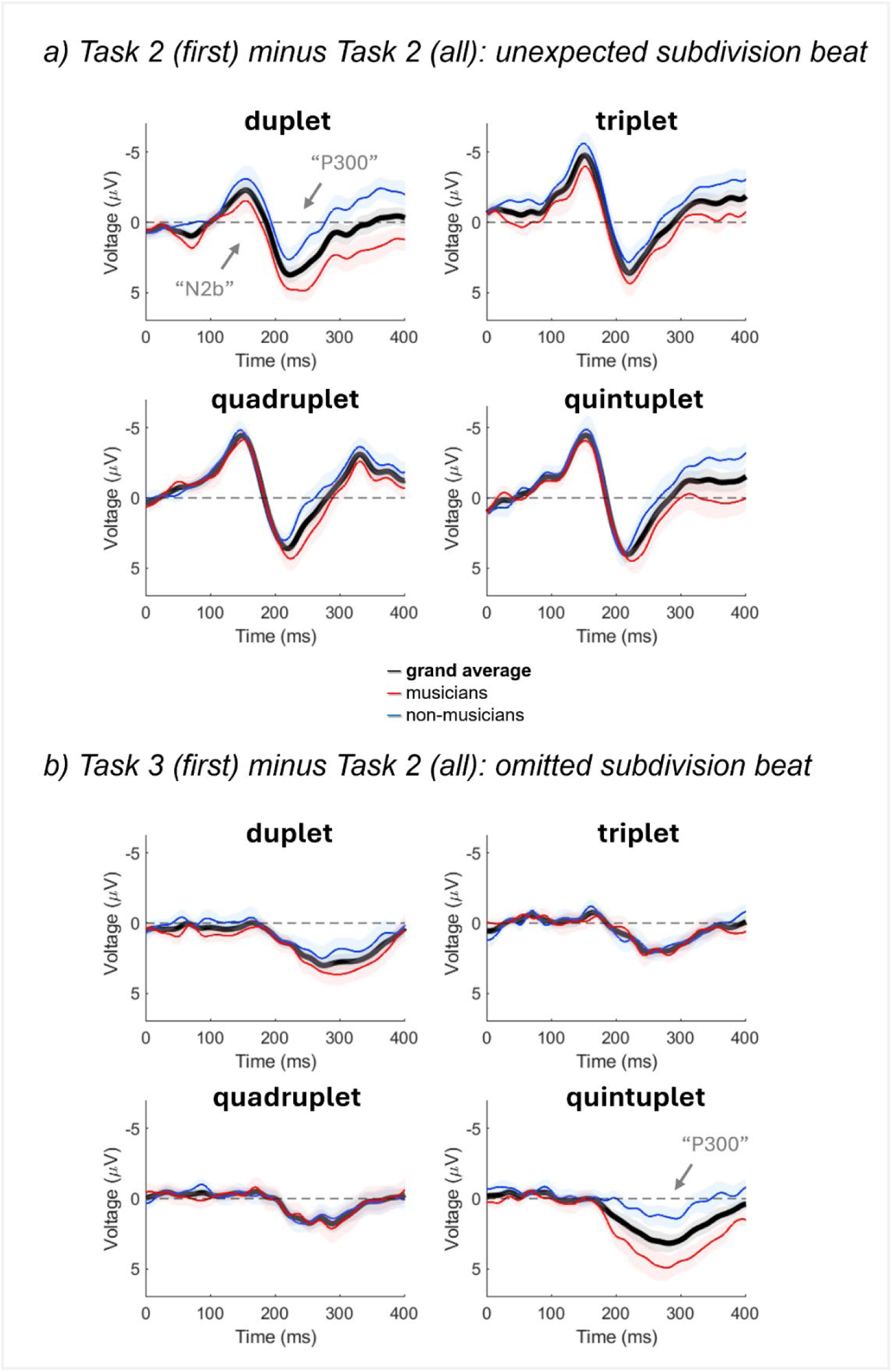
Difference waves for the first unexpected sound of each subdivision (a) or its omission (b). The difference waves were computed by subtracting the ERPs at the time of the first differentiated sound of the subdivision across all beat cycles (i.e. corresponding to second sound of all beat subdivisions in Task 2) from the ERPs at (a) the first onset subdividing the beat (Task 2 first subdivided beat cycle) or (b) its first omission (Task 3 first beat cycle). (a) At the start of Task 2, the second sound of duplets elicited smaller N2b than the second sound of other subdivisions. Musicians tended to show larger P300 deflections for the unexpected sounds of the subdivisions. (b) At the start of task 3, the P300 for the first omitted sound of quintuplets seemed to be greater in musicians. Blue and red lines respectively depict mean responses from musicians and non-musicians, while thick black lines depict participants’ grand average. Standard errors are similarly depicted around the mean lines.

We also examined expectancy violations by analyzing responses to omissions of the first subdividing sound. Here, musicians showed larger P300 responses following the omission of the first quintuplet event, suggesting that omissions within this uncommon metrical structure engaged stronger attentional updating processes. Together with the trends observed for N2b latency and P300 amplitude, these findings suggest that musical training may refine the precision and flexibility of temporal predictions, particularly for more complex rhythmic structures. Overall, our ERP results support the view that rhythm perception is naturally biased toward binary temporal organization, whereas musical expertise enhances the predictive mechanisms required to process more complex metrical structures.

### 4.3 The binary subdivision advantage and musical training

Non-musicians exhibited greater consistency in tapping to the beat of the duplets than to any other subdivision, whereas musicians showed similarly high consistency across duplets, triplets, and quadruplets. This pattern suggests that a preference for binary subdivision emerges without formal musical training, whereas musical expertise extends accurate sensorimotor synchronization to more complex metrical structures. Such expertise may arise through music-related neuroplasticity (Zatorre et al., 2007; Chen et al., 2008), which strengthens auditory-motor coupling and improves temporal predictions during SMS (Morillon and Baillet, 2017; Damm et al., 2020; Zalta et al., 2020).

Several lines of evidence suggest that the binary subdivision advantage is a fundamental property of rhythm perception rather than a consequence of musical training. A preference for binary structures can already be observed early in development, as infants more readily process duple than more complex meters (Bergeson and Trehub, 2005). Likewise, automatic binary grouping has been reported in the “tick-tock” effect (Brochard et al., 2003) and during spontaneous air-tapping to an auditory metronome (Bouvet et al., 2019). Even during spontaneous movement to an unaccented metronome, participants preferentially grouped events into binary rather than ternary units, although accenting the stimulus could shift this preference toward ternary grouping (Bouvet et al. 2020). Evidence for this binary bias also comes from learning paradigms and computational models. Non-musicians spontaneously generated binary groupings (1:2 and 2:1) when transmitting rhythmic patterns across generations in iterated learning paradigms (Ravignani et al., 2016). More recently, EEG studies identified the 1:1 and 2:1 ratios as stable neural and behavioral categories for classifying temporally jittered rhythms (Barbero et al., 2025), and similar categories emerged in computational models that transformed random temporal sequences into structured rhythmic patterns (Coissac et al, 2026).

Our findings are also consistent with previous SMS work suggesting that synchronization involves an implicit subdivision level. This subdivision level appears to emerge automatically for binary intervals (Repp, 2008b), whereas ternary subdivisions depend more strongly on rhythmic context (Repp and Jendoubi, 2009). Accordingly, binary grouping has repeatedly been observed when both musicians and non-musicians synchronize to the perceived beat in polyrhythms (Møller et al., 2021; Stupacher et al., 2025; Møller et al., 2026) or to competing melodic groupings in triple and quadruple meters (Mayayo et al., 2026). It seems, then, that temporal processes in rhythmic perception favor binary groupings of events when they are available.

### 4.4 Study limitations

One limitation of this study is that participants might have perceived subdivisions themselves as the beat, rather than maintaining the original beat throughout the sequence. This possibility is supported by the generally weak neural responses at the beat frequency (*f* in Figure 4), which reached significance only for quadruplets. In the absence of an explicit acoustic cue marking the downbeat. Without (Farrokh, 2018), participants might have anchored the perceived beat at different positions within the subdivision sequence, thereby reducing phase-locked steady-state evoked potentials after averaging across trials. Additional analyses with the fast Fourier transform before trial averaging revealed small peaks at the beat frequency for triplets and quadruplets (Figure S2), suggesting these faster events called for metrical grouping. The absence of comparable peaks for duplets and quintuplets may reflect different underlying mechanisms. For duplets, the comfortable inter-onset interval (400 ms) may have encouraged listeners to adopt the subdivision as the new beat (Damsma et al., 2025), while for quintuplets, the rapid presentation rate may have made it difficult to maintain a stable internally projected beat across successive events. Nevertheless, this limitation does not alter our principal findings: all subdivisions rhythms elicited neural synchronization at beat-related frequencies during the predictive SMS task and the ERP analyses showed smaller N2b responses for duplets than for triplets and quadruplets during the listening-only tasks, indicating differential neural processing of subdivision structure.

Another limitation concerns statistical power and group comparisons. The comparison between musicians and non-musicians was based on approximately fifteen participants per group, limiting the ability to detect modest group differences in their varying ERPs, obtained from the average of nine trials per condition. Our experimental paradigm was optimized for frequency-tagging analyses, requiring long repetitive stimulus sequences, rather than for eliciting deviant responses to events in each trial. Consequently, the group-related trends observed in the ERP measures should be interpreted cautiously. Similarly, the relationship between neural synchronization and behavioral performance appears to depend on musical expertise. Although neural responses during SMS correlated with tapping consistency when all participants were analyzed together (Table 2; see also Nozaradan, Peretz, and Keller, 2016; Gibbings et al., 2023), these correlations largely disappeared within the musician group. The ceiling effects observed for musicians’ tapping consistency likely reduced behavioral variability and therefore limited the strength of these correlations. This unexpected finding in musically trained individuals suggests that additional factors beyond overt motor performance modulate the link between performance and the frequency-tagged responses (see Damsma et al., 2025).

## 5. CONCLUSIONS

Our study investigated how the human brain synchronizes with beat subdivisions and how musical training shapes the underlying neural and behavioral responses. By asking musicians and non-musicians to synchronize their taps to the beat of four subdivision rhythms (i.e., duplets, triplets, quadruplets, and quintuplets), we found that non-musicians showed more consistent tapping for duplets than for the other subdivisions, supporting a behavioral advantage for the simplest binary (1:2) subdivision. In contrast, musicians exhibited stronger frequency-tagged responses and more consistent tapping across all common subdivision rhythms, suggesting that musical training extends efficient sensorimotor synchronization beyond binary structures. Passive listening to duplets also produced the largest beat-related neural responses and smaller N2b amplitudes than the other subdivisions, indicating that binary subdivision was more consistent with listeners’ metrical expectations and imposed lower predictive demands. These findings demonstrate that neural synchronization and predictive processing jointly contribute to the sensorimotor advantages associated with beat subdivision. Taken together, our results advance our understanding of the neural mechanisms supporting rhythm perception and sensorimotor synchronization, while highlighting the potential of binary rhythmic structures to optimize rhythmic auditory stimulation in clinical applications for individuals with movement disorders (Thaut, and Abiru, 2010), particularly those without formal musical training.

## ACKNOWLEDGEMENTS

We thank our colleagues at the Center for Music in the Brain (Aarhus) and the Center for Brain and Cognition (Barcelona) for their valuable feedback and assistance. We also thank Anne Marie Farrokh for her contributions to the design and implementation of the pilot version of this paradigm during her master’s degree.

## Author contributions

All authors had full access to the data in the study and take responsibility for the integrity of the data and the accuracy of the data analysis. Conceptualization, ACM, BK, JT, PV; Methodology, ACM, BK; Software, Investigation, Formal Analysis, ACM; Resources: PV; Writing - Original Draft, ACM; Writing - Review & Editing, ACM, BK, JT, PV; Visualization, ACM; Supervision, BK, JT, PV; Project Administration, BK; Funding Acquisition, ACM, JT, PV.

## Data and Code Accessibility Statement

The data and code that support the findings of this study are available on request from the corresponding author. The data are not publicly available due to privacy and ethical restrictions.

