## Supplementary Materials for "Neural and behavioral correlates of sensorimotor synchronization to a subdivided beat: Evidence for a binary grouping advantage and musical training effects"

#### 1. FREQUENCY-TAGGED AMPLITUDES AT EACH FREQUENCY OF INTEREST

One-sample *t*-tests against zero of the frequency-tagged amplitudes at all frequencies of interest across the four tasks and the four subdivisions of the beat.

| | Subdivision rhythm (ratio) | df | Listen to Beat | | Listen to Sub. | | Listen to Beat | | Listen to Sub. Tap to Beat | |
|---|---|---|---|---|---|---|---|---|---|---|
| | | | *t* | *p* | *t* | *p* | *t* | *p* | *t* | *p* |
| **1.25 Hz** | 1:2 | 30 | 5.201 | <.001 | 1.608 | .118 | 7.142 | <.001 | 7.067 | <.001 |
| | 1:3 | 30 | 7.236 | <.001 | .976 | .337 | 7.216 | <.001 | 6.456 | <.001 |
| | 1:4 | 30 | 5.884 | <.001 | 2.101 | .044 | 7.751 | <.001 | 6.088 | <.001 |
| | 1:5 | 30 | 4.956 | <.001 | -.496 | .624 | 8.891 | <.001 | 3.427 | .002 |
| **2.5 Hz** | 1:2 | 30 | 5.568 | <.001 | 10.359 | <.001 | 5.859 | <.001 | 9.827 | <.001 |
| | 1:3 | 30 | 6.217 | <.001 | .922 | .364 | 7.776 | <.001 | 5.295 | <.001 |
| | 1:4 | 30 | 7.095 | <.001 | 2.326 | .027 | 8.707 | <.001 | 5.299 | <.001 |
| | 1:5 | 30 | 5.846 | <.001 | .443 | .661 | 8.374 | <.001 | 1.739 | .092 |
| **3.75 Hz** | 1:2 | 30 | 6.638 | <.001 | -1.149 | .260 | 6.997 | <.001 | 5.701 | <.001 |
| | 1:3 | 30 | 6.343 | <.001 | 8.564 | <.001 | 5.745 | <.001 | 9.996 | <.001 |
| | 1:4 | 30 | 6.011 | <.001 | -.515 | .610 | 5.305 | <.001 | 5.470 | <.001 |
| | 1:5 | 30 | 7.052 | <.001 | .210 | .835 | 5.350 | <.001 | 1.431 | .163 |
| **5 Hz** | 1:2 | 30 | 9.193 | <.001 | 9.950 | <.001 | 7.307 | <.001 | 8.159 | <.001 |
| | 1:3 | 30 | 7.526 | <.001 | .882 | .385 | 6.882 | <.001 | 4.171 | <.001 |
| | 1:4 | 30 | 7.123 | <.001 | 10.629 | <.001 | 5.530 | <.001 | 10.36 | <.001 |
| | 1:5 | 30 | 8.467 | <.001 | -.744 | .463 | 5.036 | <.001 | 1.840 | .076 |
| **6.25 Hz** | 1:2 | 30 | 8.680 | <.001 | 1.183 | .246 | 6.006 | <.001 | 2.796 | .009 |
| | 1:3 | 30 | 7.106 | <.001 | .036 | .972 | 6.434 | <.001 | 2.762 | .010 |
| | 1:4 | 30 | 7.246 | <.001 | -.109 | .914 | 4.898 | <.001 | 3.188 | .003 |
| | 1:5 | 30 | 6.223 | <.001 | 9.262 | <.001 | 5.074 | <.001 | 9.214 | <.001 |

**Table S1. Results of statistical analyses performed to detect corrected neural amplitudes that differ from zero.** One-sample t-tests were conducted at each frequency of interest and subdivision rhythm, including duplets, 1:2, triplets, 1:3, quadruplets, 1:4, and quintuplets, across four different tasks: listening to the beat, listening to the subdivision, listening to the beat while tapping, and listening to the subdivision while tapping. The table reports the *t*-statistic and *p*-value for each normalized neural amplitude. The rows where the frequency of interest matches the frequency of the subdivision rhythm are highlighted in gray. The abbreviation "df" stands for degrees of freedom. The results show that for all four subdivisions, the t-tests against zero revealed the presence of SSEPs at all frequencies during the listen-to-the-beat tasks (all $p < .001$). Similarly, during the listen-to-the-subdivision and tapping-to-the-beat tasks, SSEPs were elicited at the frequencies related to each subdivision (all $p < .001$). Notably, a peak at 1.25 Hz was observed for quadruplets during the listen-to-the-subdivision task ($p < .044$) and for all subdivisions during the tapping-to-the-beat task (all $p \leq .002$). When applying the Bonferroni correction to the sixteen tests comprised of each frequency, the level of significance drops to a conservative $p = .003$. Nonetheless, the *p*-values for the main peaks reported here remained above this level. Thus, the analysis identified several peaks in the frequency spectra of the beat and its four subdivisions, centered at 1.25, 2.5, 3.75, 5, and 6.25 Hz (see Figure 2).

FIGURE S1: EEG AMPLITUDES ACROSS TASKS AND GROUPS

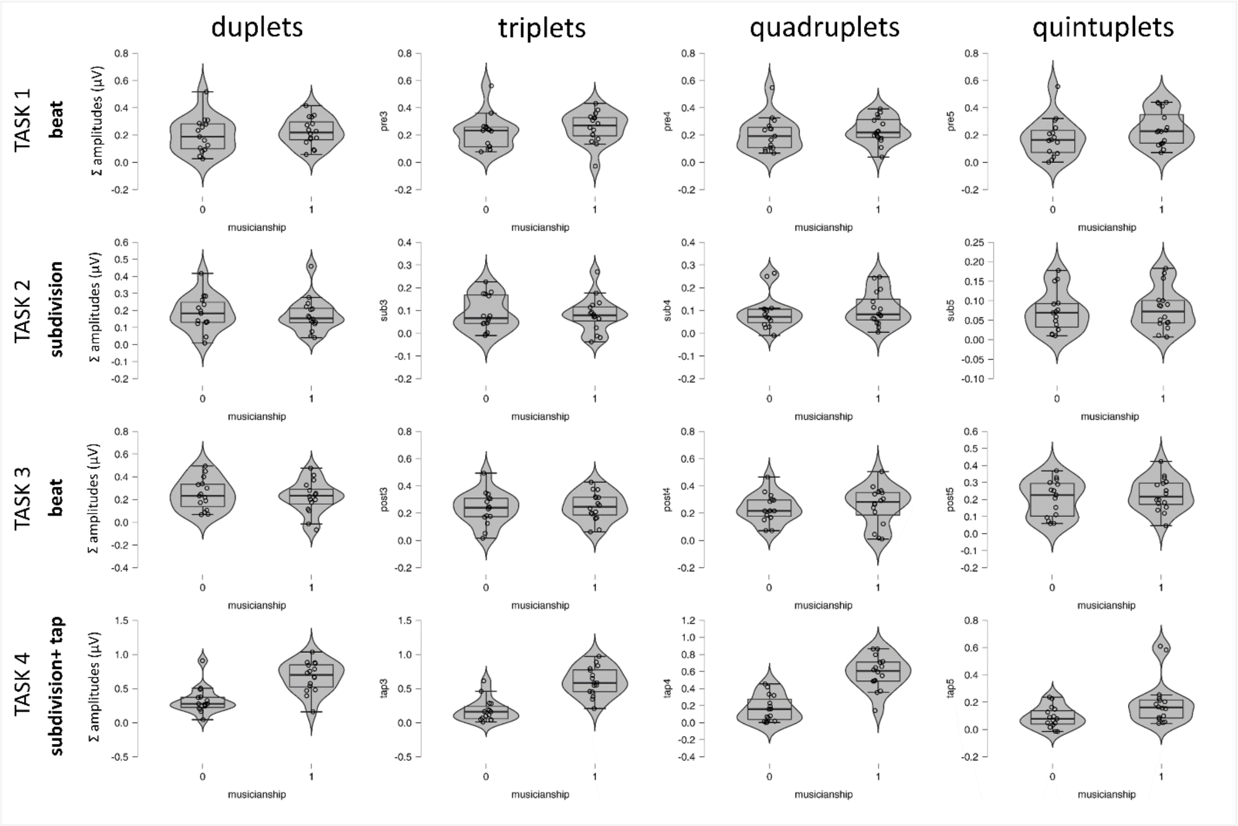


**Figure S1. Scatter box violin plots of the summated noise-corrected amplitudes.** The frequency-tagged amplitudes at the five frequencies of interest were summated. Non-musicians labeled as “0” and musicians labeled as “1”. This figure aims to depict the processed EEG data combining different visualizations.

#### 2. FREQUENCY-TAGGED AMPLITUDES AVERAGING TRIALS AFTER THE FFT

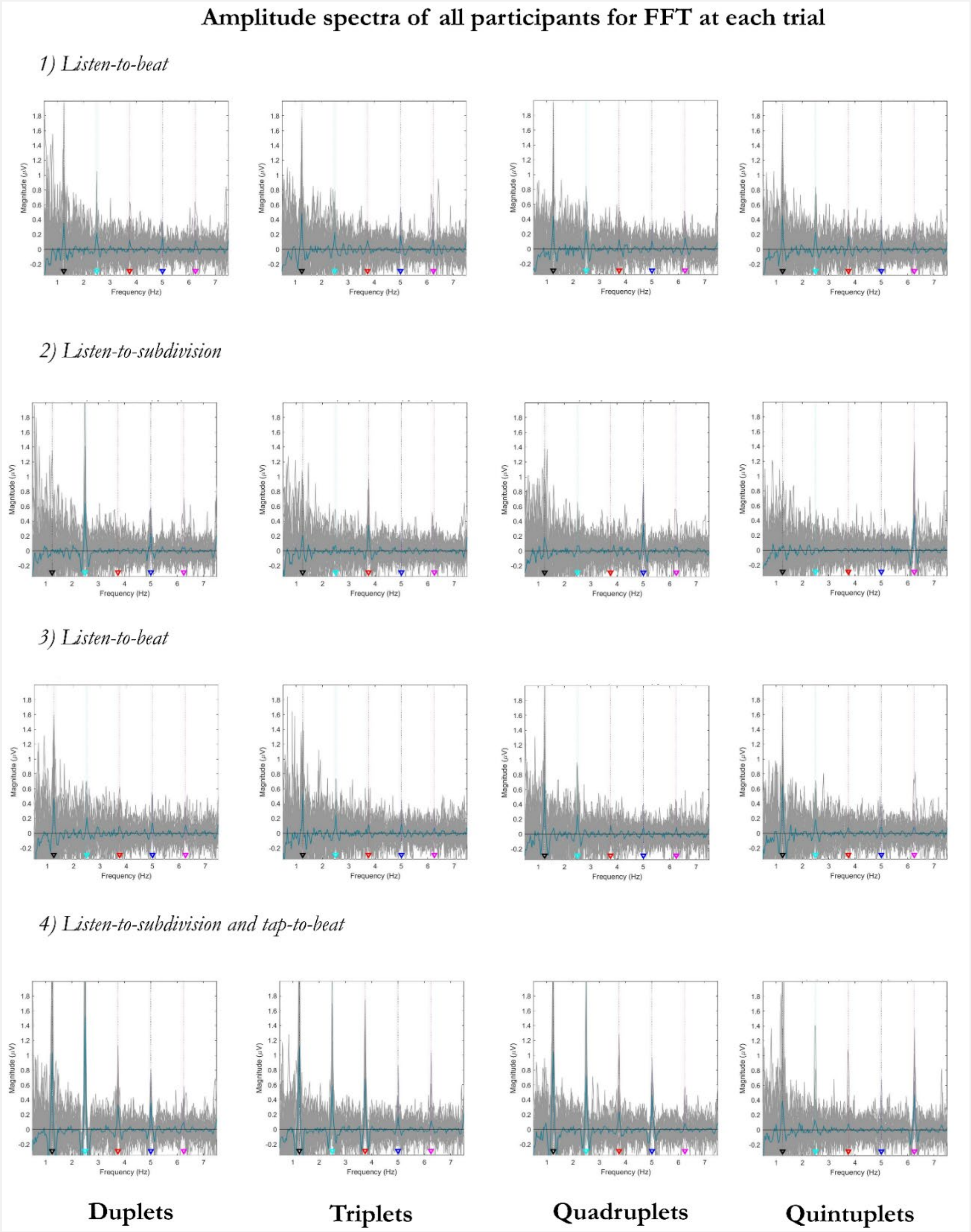


**Figure S2**. **Frequency spectra obtained for each task and subdivision after applying the Fast Fourier transform before averaging the trials.** The gray lines represent the amplitudes of all participants, while the black line represents the mean amplitude for each task (rows) and subdivision (columns). The triangles highlight the frequencies of the beat and its harmonics: black for the beat frequency (*f*), cyan for the duplet frequency (2*f*), red for the triplet frequency (3*f*), blue for the quadruplet frequency (4*f*), and pink for the quintuplet frequency (5*f*). Note in (2) small peaks at the beat frequency for triplets and quadruplets.

#### 3. EVENT-RELATED POTENTIALS AND DIFFERENCE WAVES ANALYSES

**Table S2. Results of one-sample *t*-tests against zero for the positive and negative peaks in the difference waves elicited by each subdivision sound onset detection or omission.** The table reports the mean, standard deviation, *t*-statistic and *p*-value for each difference wave mean peak amplitudes (averaging amplitudes within a time window of 60 ms centered at the global latency across participants). These amplitudes reflect the unexpected sound onset of a beat subdivision (i.e., Task 2-first minus Task 2-all) or the omission of an expected sound onset (i.e., Task 3-first minus Task 2-all). The average of all beat cycles from task 2 is used as a baseline (or standard) to compute the difference waves and reveal mismatch responses. Degrees of freedom are 30, and significance level is not corrected for multiple comparisons. Gray background highlights non-significant peaks.

| | | **Negative mean peak amplitudes** (N2b between 50-180ms) | | **Positive mean peak amplitudes** (P300 between 180-300ms) | |
|---|---|---|---|---|---|
| | | Mean (SD) | *t (p)* | Mean (SD) | *t (p)* |
| **Unexpected sound onset (Task 2)** | Duplet | -1.627 (2.955) | -3.065 (.005) | 3.052 (2.830) | 6.004 (<.001) |
| | Triplet | -3.544 (3.285) | -6.006 (<.001) | 2.639 (3.307) | 4.443 (<.001) |
| | Quadruplet | -3.394 (2.335) | -8.092 (<.001) | 2.702 (2.696) | 5.580 (<.001) |
| | Quintuplet | -3.278 (2.901) | -6.290 (<.001) | 3.039 (2.981) | 5.677 (<.001) |
| **Unexpected sound omission (Task 3)** | Duplet | 0.240 (2.187) | 0.612 (.545) | 2.771 (2.957) | 5.218 (<.001) |
| | Triplet | -0.335 (1.977) | -0.943 (.353) | 1.739 (2.428) | 3.987 (<.001) |
| | Quadruplet | -0.304 (2.439) | -0.693 (.493) | 1.459 (3.211) | 2.530 (.017) |
| | Quintuplet | -0.356(1.219) | -1.626 (.114) | 3.008 (3.441) | 4.868 (<.001) |

**Figure S3. Event-related potentials obtained for each subdivision during the first beat cycle of each task (a), the last beat cycle of each task (b) and averaging all the beat cycles of each task (c).** ERPs are depicted for 800 ms, the length of the beat cycle, after applying an inside-epoch baseline from 0 to 100 ms to center participants' epochs around zero before averaging them across trials. For each task (columns) and subdivision (rows), color lines represent the amplitudes of each participant, while the black line represents the mean amplitude. Vertical gray dotted lines mark the onset of each subdivision sound onsets. The red arrows point to N1-P2 responses elicited by the (unexpected) first sound of each subdivision, while the blue arrows point to positive neural responses (P300) elicited by the omission of the expected first sound onset of each subdivision. The yellow arrows point to the ERPs used as a baseline to calculate the N2b and P300 in the difference waves.

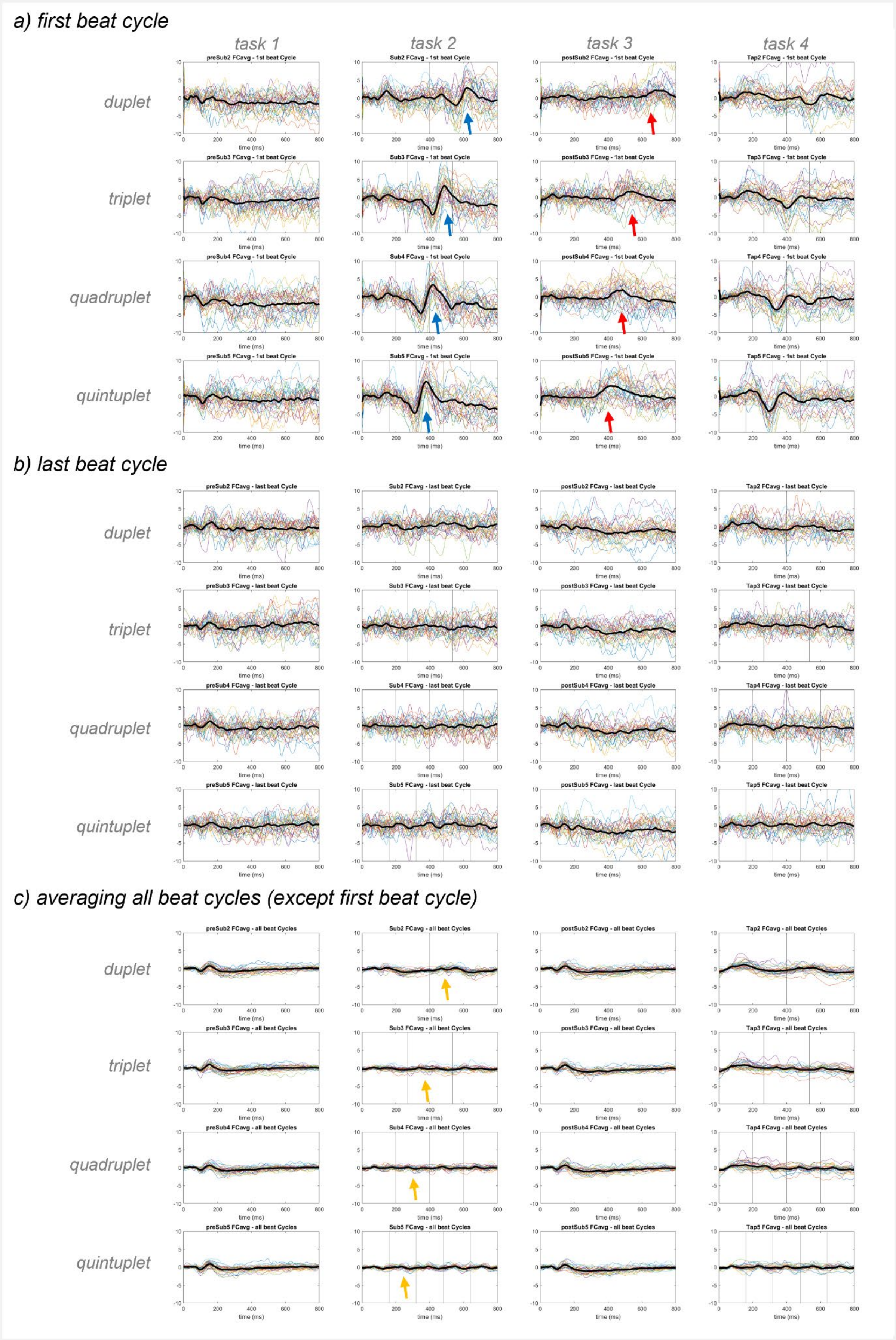
a) first beat cycle
task 1
task 2
task 3
task 4
duplet
triplet
quadruplet
quintuplet
preSub2 FCavg - 1st beat Cycle
Sub2 FCavg - 1st beat Cycle
postSub2 FCavg - 1st beat Cycle
Tap2 FCavg - 1st beat Cycle
preSub3 FCavg - 1st beat Cycle
Sub3 FCavg - 1st beat Cycle
postSub3 FCavg - 1st beat Cycle
Tap3 FCavg - 1st beat Cycle
preSub4 FCavg - 1st beat Cycle
Sub4 FCavg - 1st beat Cycle
postSub4 FCavg - 1st beat Cycle
Tap4 FCavg - 1st beat Cycle
preSub5 FCavg - 1st beat Cycle
Sub5 FCavg - 1st beat Cycle
postSub5 FCavg - 1st beat Cycle
Tap5 FCavg - 1st beat Cycle
b) last beat cycle
preSub2 FCavg - last beat Cycle
Sub2 FCavg - last beat Cycle
postSub2 FCavg - last beat Cycle
Tap2 FCavg - last beat Cycle
preSub3 FCavg - last beat Cycle
Sub3 FCavg - last beat Cycle
postSub3 FCavg - last beat Cycle
Tap3 FCavg - last beat Cycle
preSub4 FCavg - last beat Cycle
Sub4 FCavg - last beat Cycle
postSub4 FCavg - last beat Cycle
Tap4 FCavg - last beat Cycle
preSub5 FCavg - last beat Cycle
Sub5 FCavg - last beat Cycle
postSub5 FCavg - last beat Cycle
Tap5 FCavg - last beat Cycle
c) averaging all beat cycles (except first beat cycle)
preSub2 FCavg - all beat Cycles
Sub2 FCavg - all beat Cycles
postSub2 FCavg - all beat Cycles
Tap2 FCavg - all beat Cycles
preSub3 FCavg - all beat Cycles
Sub3 FCavg - all beat Cycles
postSub3 FCavg - all beat Cycles
Tap3 FCavg - all beat Cycles
preSub4 FCavg - all beat Cycles
Sub4 FCavg - all beat Cycles
postSub4 FCavg - all beat Cycles
Tap4 FCavg - all beat Cycles
preSub5 FCavg - all beat Cycles
Sub5 FCavg - all beat Cycles
postSub5 FCavg - all beat Cycles
Tap5 FCavg - all beat Cycles
time (ms)

#### 4. ANALYSES ON EACH FREQUENCY OF INTEREST ACROSS TASKS AND GROUPS

***Exploratory research questions (RQ) and statistical tests to answer them***

*RQ1:* The first RQ asked *how the brain responds to the beat after listening to different subdivisions.* We compared the effect of listening to distinct subdivisions at each frequency of interest (1.25, 2.5, 3.75, 5, 6.25 Hz). A three-way mixed design ANOVA was conducted for the peak magnitudes of each frequency of interest, using the within-factors Task (Pre-Subdivision, Post-Subdivision) and Rhythm (Duplets, Triplets, Quadruplets, Quintuplets), and the between-factor Group (Musicians, non-Musicians).

*RQ2:* The second RQ explored changes in *brain responses when people passively listen to subdivisions compared to when they listen and simultaneously tap the beat.* To answer this question, we ran three-way mixed design ANOVAs again for each frequency of interest using the within-factor Task (Subdivision, Tapping).

*RQ3*: The third RQ examined *significant changes across frequencies in RQ1 and RQ2*. To address this question for RQ1, we subtracted the Pre-Subdivision amplitudes from those of the Post-Subdivision, and for RQ2, we subtracted the Subdivision amplitudes from those of the Tapping. The resulting "values" were analyzed in two separate three-way mixed design ANOVAs, using the within-factors Frequency (1.25, 2.5, 3.75, 5, 6.25 Hz) and Rhythm (Duplets, Triplets, Quadruplets, Quintuplets), and the between-factor Group (Musicians, non-Musicians), followed by corrected post hoc comparisons.

We corrected the degrees of freedom using Greenhouse-Geisser estimates of sphericity when the assumptions for sphericity were violated. When significant effects and/or interactions appeared, we conducted post hoc two-tailed t-tests with the Bonferroni correction (with significance level at $p < .05$) for all analyses.

##### *Results of exploratory research question 1*

Results pertaining to RQ1 (“*How does the brain respond to the beat after listening to different subdivisions?”)* showed a significant effect of Task on the peak magnitudes centered at 1.25, 2.5 and 6.25 Hz (all $p \leq .021$, see Table S3a). The *post hoc* pairwise comparisons revealed that the peaks at 1.25 Hz ($M_{pre}$ = 0.09, SE = .014, $M_{post}$ = 0.11, SE = .012, MD = 0.02, $p = .021$) and 2.5 Hz ($M_{pre}$ = 0.04, SE = .006, $M_{post}$ = 0.05, SE = .006, MD = 0.01, $p = .019$) increased in amplitude after listening to subdivisions. In contrast, the amplitudes at 6.25 Hz decreased after listening to the subdivisions ($M_{pre}$ = 0.03, SE = .003, $M_{post}$ = 0.02, SE = .003, MD = 0.01, $p = .018$). The absence of any interaction may indicate that these increases of the peaks of the beat and its first harmonic occurred after listening to all subdivision rhythms in both musicians and non-musicians (see Figure S2).

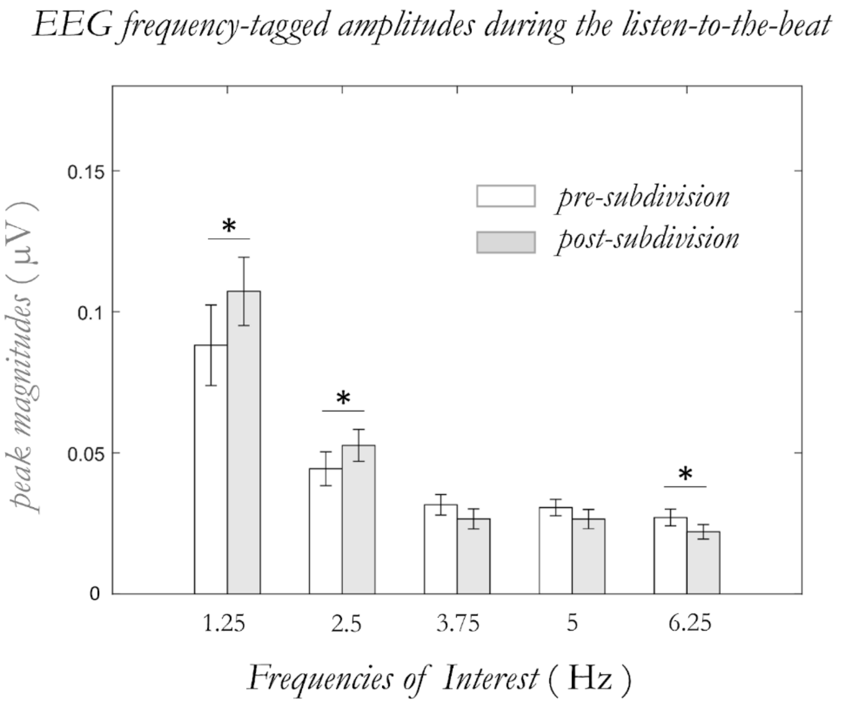


**Figure S4. Peak amplitudes at each frequency of interest during the listening-to-the-beat tasks (tasks 1 and 3)**. The bars represent the amplitude values before (in white) and after (in gray) listening to the subdivisions. An asterisk (*) indicates statistical significance at $p < .05$. These results suggest that there was an increase in amplitude at the beat and its first harmonic after listening to subdivisions, which is relevant to the RQ1 being investigated.

### TABLE S3

**a) Frequency-tagged amplitudes from *Listening to beat before subdivisions* versus *Listening to beat after subdivisions***

| FoI | | Task | | | | Rhythm | | | | Task*Rhythm | | | | Group | | | |
|---|---|---|---|---|---|---|---|---|---|---|---|---|---|---|---|---|---|
| | | df | F | *p* | $\eta^2$ | df | F | *p* | $\eta^2$ | df | F | *p* | $\eta^2$ | df | F | *p* | $\eta^2$ |
| *1.25 Hz* | | 1,29 | 5.93 | **.021** | .170 | 2.3,67.5 | 1.43 | .246 | .047 | 3,87 | 1,28 | .285 | .042 | 1,29 | 0.26 | .613 | .009 |
| | *Group | 1,29 | 0.14 | .708 | .005 | 2.3,67.5 | 0.64 | .555 | .022 | 3,87 | 0.48 | ,695 | .016 | | | | |
| *2.5 Hz* | | 1,29 | 6.18 | **.019** | .176 | 3,87 | 1.05 | .375 | .035 | 2.4,70.6 | 0.15 | .896 | .005 | 1,29 | 0.02 | .901 | .001 |
| | *Group | 1,29 | 0.63 | .433 | .021 | 3,87 | 0.92 | .434 | .031 | 2.4,70.6 | 2.33 | .094 | .074 | | | | |
| *3.75 Hz* | | 1,29 | 3.03 | .093 | .094 | 3,87 | 0.48 | .697 | .016 | 3,87 | 1.32 | .272 | .044 | 1,29 | 1.08 | .308 | .036 |
| | *Group | 1,29 | 1.19 | .285 | .039 | 3,87 | 0.55 | .650 | .019 | 3,87 | 0.27 | .851 | .009 | | | | |
| *5 Hz* | | 1,29 | 2.18 | .151 | .070 | 3,87 | 2.99 | **.035** | .094 | 3,87 | 1.16 | .332 | .038 | 1,29 | 0.06 | .806 | .002 |
| | *Group | 1,29 | 1.30 | .263 | .043 | 3,87 | 1.27 | .291 | .042 | 3,87 | 0.66 | .582 | .022 | | | | |
| *6.25 Hz* | | 1,29 | 6.24 | **.018** | .177 | 3,87 | 1.45 | .234 | .048 | 3,87 | 0.29 | .833 | .010 | 1,29 | 0.00 | .984 | .000 |
| | *Group | 1,29 | 2.99 | .094 | .094 | 3,87 | 1.49 | .224 | .049 | 3,87 | 1.71 | .170 | .056 | | | | |

**b) Frequency-tagged amplitudes from *Listening to subdivision* versus *Listening to subdivision while tapping to the beat***

| FoI | | Task | | | | Rhythm | | | | Task*Rhythm | | | | Group | | | |
|---|---|---|---|---|---|---|---|---|---|---|---|---|---|---|---|---|---|
| | | df | F | *p* | $\eta^2$ | df | F | *p* | $\eta^2$ | df | F | *p* | $\eta^2$ | df | F | *p* | $\eta^2$ |
| *1.25 Hz* | | 1,29 | 58.97 | **<.001** | .670 | 2.3,68.1 | 15.84 | **<.001** | .353 | 2.3,65.9 | 13.87 | **<.001** | .324 | 1,29 | 15.15 | **.001** | .343 |
| | *Group | 1,29 | 15.05 | **.001** | .342 | 2.3,68.1 | 5.33 | **.005** | .155 | 2.3,65.9 | 5.44 | **.005** | .158 | | | | |
| *2.5 Hz* | | 1,29 | 41.60 | **<.001** | .589 | 1.9,55.1 | 91.90 | **<.001** | .760 | 2.2,63.2 | 17.97 | **<.001** | .383 | 1,29 | 20.7 | **<.001** | .416 |
| | *Group | 1,29 | 24.51 | **<.001** | .458 | 1.9,55.1 | 3.28 | **.047** | .102 | 2.2,63.2 | 9.59 | **<.001** | .249 | | | | |
| *3.75 Hz* | | 1,29 | 40.88 | **<.001** | .585 | 1.8,51.0 | 81.55 | **<.001** | .738 | 3,87 | 8.66 | **<.001** | .230 | 1,29 | 14.08 | **.001** | .327 |
| | *Group | 1,29 | 21.37 | **<.001** | .424 | 1.8,51.0 | 1.51 | .233 | .049 | 3,87 | 1.50 | .221 | .049 | | | | |
| *5 Hz* | | 1,29 | 14.52 | **.001** | .334 | 1.9,55.7 | 72.68 | **<.001** | .715 | 2.1,61.2 | 1.58 | .213 | .052 | 1,29 | 12.84 | **.001** | .307 |
| | *Group | 1,29 | 9.70 | **.004** | .251 | 1.9,55.7 | 1.62 | .209 | .053 | 2.1,61.2 | 1.17 | .320 | .039 | | | | |
| *6.25 Hz* | | 1,29 | 5.16 | **.031** | .151 | 1.6,45.1 | 53.28 | **<.001** | .648 | 2.5,71.6 | 3.43 | **.029** | .106 | 1,29 | 3.66 | .066 | .112 |
| | *Group | 1,29 | 9.43 | **.005** | .245 | 1.6,45.1 | 0.31 | .681 | .011 | 2.5,71.6 | 2.36 | .090 | .075 | | | | |

**c) Difference in frequency-tagged amplitudes between tasks with the same auditory rhythms**

| *subtraction* | | Frequency | | | | Rhythm | | | | Frequency*Rhythm | | | | Group | | | |
|---|---|---|---|---|---|---|---|---|---|---|---|---|---|---|---|---|---|
| | | df | F | *p* | $\eta^2$ | df | F | *p* | $\eta^2$ | df | F | *p* | $\eta^2$ | df | F | *p* | $\eta^2$ |
| *Post-Pre* | | 1.7,50.2 | 7.53 | **.002** | .206 | 3,87 | 0.55 | .649 | .019 | 6.6,190 | 1.07 | .382 | .036 | 1,29 | 1.51 | .230 | .049 |
| | *Group | 1.7,50.2 | 0.01 | .991 | .000 | 3,87 | 0.38 | .770 | .013 | 6.6,190 | 1.11 | .359 | .037 | | | | |
| *Tap-Sub* | | 2.0,58.7 | 28.80 | **<.001** | .498 | 2.1,61.3 | 32.16 | **<.001** | .526 | 5.5,158.2 | 6.33 | **<.001** | .179 | 1,29 | 38.89 | **<.001** | .573 |
| | *Group | 2.0,58.7 | 7.78 | **.001** | .212 | 2.1,61.3 | 12.87 | **<.001** | .307 | 5.5,158.2 | 3.14 | **.008** | .098 | | | | |

**Table S3. Main effects and interactions of EEG frequency-tagged amplitudes.** Results of the 3-way Mixed Design ANOVAs performed to assess the main effects and interactions of EEG frequency-tagged amplitudes. The table provides information on the degrees of freedom, F-statistic, *p*-value, and $\eta^2$ effect size for three sets of comparisons. In (a), the ANOVA examined the amplitudes from listening to the beat before and after listening to each distinct subdivision, at five frequencies of interest (FoI). In (b), the ANOVA investigated the amplitudes from listening to four subdivisions passively versus simultaneously tapping to the beat, at five frequencies of interest. In (c), the ANOVA assessed the differences in amplitudes between listening-to-the-beat or listening-to-the-subdivision tasks across five frequencies and four subdivision rhythms. Horizontally, (a) and (b) show the within factors *Task* and *Rhythm*, the interaction *Task*Rhythm*, and the between factor Group (separating musicians and non-musicians). In (c), the within-factors *Frequency* and *Rhythm*, the interaction *Frequency*Rhythm*, and the between factor *Group* are presented. Additionally, all the interactions involving the between factor *Group* are reported in another row below the within-factor main effects and interaction. Significant *p*-values are indicated in bold.

##### *Results of exploratory research question 2*

Results pertaining to RQ2 ("*What are the changes in brain responses when people passively listen to subdivisions compared to when they listen and simultaneously tap the beat?*") revealed multiple main effects and interactions across the five frequencies (see Table S3b). Importantly, we found a significant interaction between Task and Group across all frequencies (all $p \leq .005$), as well as between Task and Rhythm (all $p \leq .029$, except at 5 Hz). Furthermore, Task, Group and Rhythm significantly interacted with each other at 1.25 and 2.5 Hz (all $p \leq .005$). The *post hoc* pairwise comparisons revealed three main findings (see Figure S3). First, musicians' amplitudes were greater than non-musicians' amplitudes in the tapping task (see Table S4). This was true for all subdivisions at 3.75, 5 and 6.25 Hz, for triplets and quadruplets at 1.25 and 2.5 Hz, and for duplets at 2.5 Hz. Second, the amplitudes in the tapping task were generally greater than the amplitudes in the listening task (see Table S4 and Table S5). This effect was mainly found in musicians across all subdivisions and frequencies (except for quintuplets at 2.5 Hz, $p > .05$), and only found in non-musicians for duplets and triplets at 1.25 Hz (see Table S4). Besides, the increase due to tapping became non-significant for quintuplets at 3.75 and 6.25 Hz, and for duplets at 6.25 Hz (see Table S5). Third, the amplitudes at the frequency of each subdivision were higher than the amplitudes at frequencies unrelated to the subdivision for both musicians and non-musicians during listening and tapping (see Table S6). Interestingly, the amplitudes related to quintuplets were normally smaller: at 4 Hz for all participants during both tasks, at 3.75 Hz in the tapping task, and at 1.25 and 2.5 Hz in the tapping task (but only in musicians). Finally, at 1.25 Hz, non-musicians had higher amplitudes when they tapped to the beat of duplets, compared to quadruplets and quintuplets.

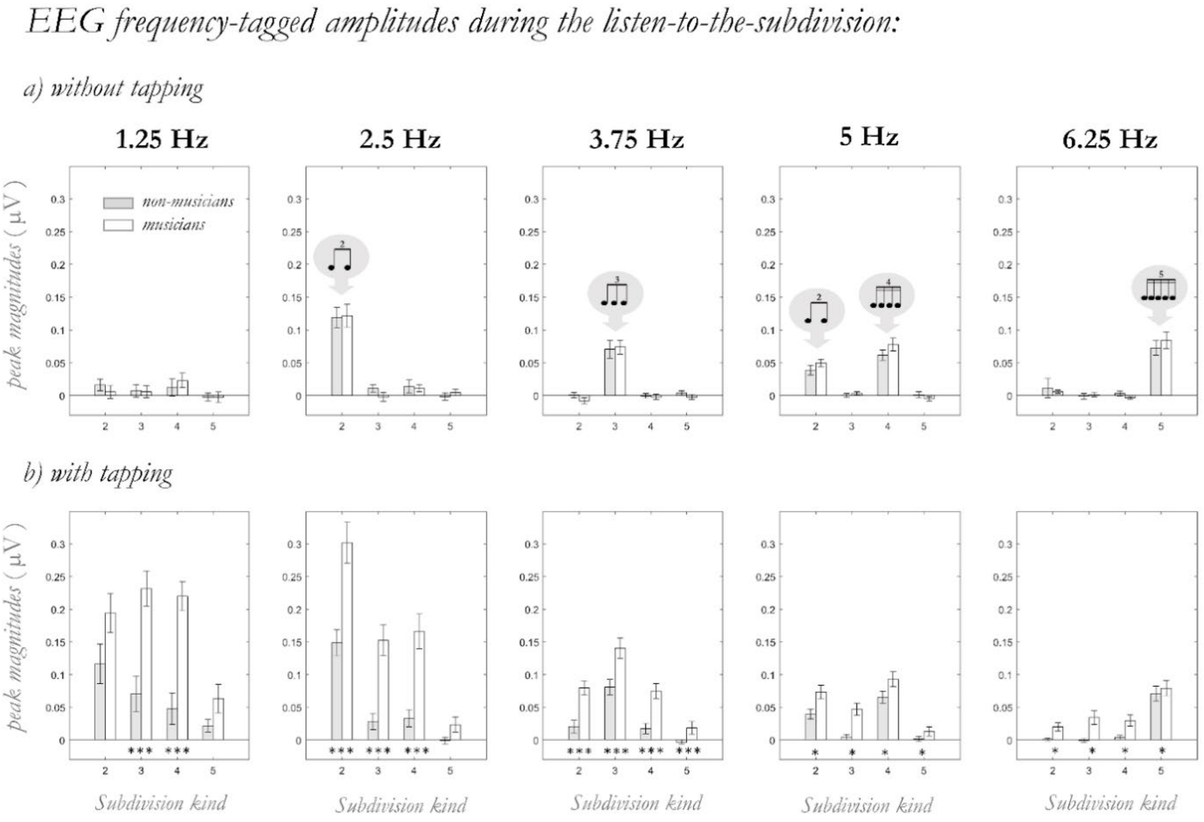


**Figure S5. Peak amplitudes at each frequency of interest and subdivision during the listening to the subdivisions (a) without tapping (task 2) and (b) with tapping to the beat (task 4).** The bars depict musicians (in white) and non-musicians (in gray). An asterisk (*) indicates statistical significance $p < .05$; while three asterisks (***) indicate $p < .001$. Regarding RQ2, the results show a general increase in amplitudes when participants tap the beat. Furthermore, it is notable that musicians had higher amplitudes than non-musicians in almost all beat-related frequencies and subdivision types.

##### *Results of exploratory research question 3*

Results pertaining to RQ3 (“*Are there significant changes across frequencies in RQ1 and RQ2?*”) showed a main effect of Frequency (see Table S3c). The *post hoc* pairwise comparisons (depicted in Figure S4a) indicated that the increases at 1.25 Hz (M = .019, SE = 008) were greater than the increases at 5 Hz (M = -.004, SE = .003, MD = .023, $p$ = .023), and that the increases at 2.5 Hz (M = .008, SE = 003) were greater than the increases at 3.75 Hz (M = -.005, SE = .003, MD = .013, $p$ = .021), 5 Hz (M = -.004, SE = .003, MD = .012, $p$ = .014), and 6.25 Hz (M = -.005, SE = .002, MD = .013, $p$ = .025). For the Tap-Sub subtraction, the ANOVA revealed a significant interaction between Frequency, Rhythm and Group (see Table S3c). The *post hoc* pairwise comparisons (depicted in Figure S4b) showed that, in non-musicians, the increases at 1.25 Hz were larger for duplets than in quadruplets ($p$ = .015), while in musicians, the increases at 1.25, 2.5 and 3.75 Hz were larger for duplets, triplets and quadruplets compared to quintuplets (all $p \leq$ .038), the increases at 5 Hz were larger for triplets compared to quadruplets and quintuplets (all $p \leq$ .033) and the increases at 6.25 Hz were smaller for quintuplets compared to triplets and quadruplets (all $p \leq$ .013).

*EEG frequency-tagged amplitude differences (a) after listening to subdivisions and (b) when tapping to the beat of the subdivisions*

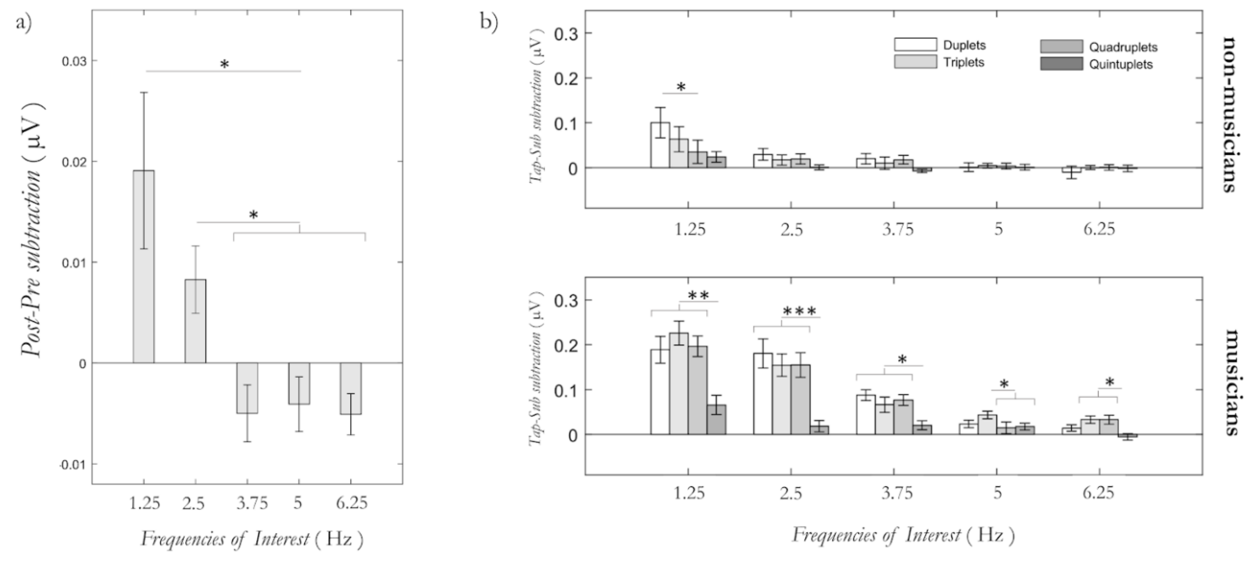

**Figure S6.** Amplitude increases across frequencies. Amplitude differences (a) between Task3-Task1, when participants listened to the beat, and (b) between Task4-Task2, when non-musicians (first row) and musicians (second row) listened to the distinct subdivisions. An asterisk (*) indicates statistical significance at $p < .05$, two asterisks (**) indicate $p < .01$ and three asterisks (***) indicate $p < .001$. Regarding RQ3, the findings in (a) indicate that the amplitude increases were greater at the frequency of the beat and its first harmonic compared to other frequencies. In (b), the results suggest that the increases in amplitude across frequencies were generally similar among subdivision types in non-musicians, while they were greater for duplets, triplets, and quadruplets in musicians.

After observing an increase in amplitude at the frequency of the beat and its first harmonic following exposure to all four subdivisions, we explored whether this increase might be related to better tapping performance. We conducted Spearman's rank correlations between the consistency of tapping to each subdivision rhythm and the increases in EEG amplitudes (post – pre listening to subdivision) at 1.25 and 2.5 Hz. However, we found no significant correlations (all $p > .05$). These findings suggest that increased entrainment to the beat during the listening tasks does not necessarily result in better tapping responses.

##### *Brief discussion on the results of the exploratory research questions*

As in the main manuscript, this extended analysis demonstrates that Sensorimotor Synchronization (SMS) can enhance neural synchronization to the beat and its harmonics. However, the effects of SMS vary at each frequency of interest, depending on factors such as musical training and the number of subdivisions. These findings suggest that the brain's ability to entrain to rhythmic stimuli is a complex process that is modulated by various factors.

###### *Neural synchronization increases during the tapping task*

Consistent with the main analyses, in this extended analysis, we found that musicians' and non-musicians' frequency-tagged amplitudes differed only during the sensorimotor task, specifically at the beat frequency, and not across all subdivisions. Additionally, we did not find group differences between the two groups when tapping the beat of quintuplets, suggesting a general disadvantage in SMS likely due to the scarcity of quintuplets in Western music or the fast tempo rate of their inter-onset intervals (IOIs). Interestingly, there were no group differences in neural synchronization to the beat of duplets (at 1.25 Hz), despite musicians being more accurate during tapping. This finding could indicate a default advantage for SMS to duplets, visible in non-musicians, but overshadowed in highly trained musicians due to their extensive practice of triplets and quadruplets. These results further highlight the complex interplay between musical training and the brain's ability to entrain to rhythmic stimuli, which is shaped by various factors such as the type of subdivision and the tempo rate.

In addition to the main findings, there were other interesting differences observed between the listening-to-the-subdivision task and the tapping-to-the-beat task. Specifically, musicians exhibited clear increases in neural amplitudes across frequencies of interest and subdivisions, while non-musicians only showed increases at the frequency of the beat for triplets and quadruplets. When tapping to the beat of the subdivision, musicians displayed increased amplitudes across frequencies and subdivision rhythms (Figure S2b), except for the quintuplets (Figure S3b). These results suggest that increased subdivision may improve temporal predictions for musicians. However, the amplitudes decreased at higher frequencies and faster rhythms. In contrast, non-musicians exhibited neural increases only for duplets and triplets at the frequency of the beat (1.25 Hz). Notably, these subdivisions correspond to the smallest integer ratios found across cultures (Savage et al., 2015) and are commonly reproduced by both adult and child non-musicians (Drake, 1998).

These findings suggest that the brain's ability to synchronize with rhythmic stimuli can be modulated by extended training, while it is shaped by cultural background and biological factors.

*Neural synchronization increases after listening to the subdivisions*

Another relevant finding of the extended analyses of the study is that after listening to different subdivision rhythms, the frequency-tagged peaks increased at the frequency of the beat and its first harmonic. This increase could reflect a stabilization of neural synchronization to the frequency of the beat and its simplest subdivision (1:2 ratio), which is consistent with the small integer ratio bias in timing (discussed in Ravignani et al., 2018; also see Mathias et al., 2020) and could explain the presence of duplets in all musical idioms (Savage et al., 2015).

Alternatively, these increases could reflect a general attentional gain to prepare for tapping to the beat, as well as cognitive processes that enhance the beat based on its simplest binary subdivision. Notably, this preparation does not seem to reflect covert movement to the beat, as the topographies in Figure 1 do not show left-lateralized activity related to the contralateral right hand, as seen in the tapping condition. Similarly, the lack of correlations between tapping performance and these neural increases also suggest that no covert movement was performed before the SMS task. In line with attentional gains, stronger activity for the first harmonic compared to the beat frequency during SMS has been reported in the bilateral middle/posterior cingulate gyrus, an area involved in attention-related motor control (Pretto, Deiber, & James, 2018; Vogt, 2009; Leech & Sharp, 2014). However, this difference disappeared in passive listening. Tierney and Kraus (2014) also connected increases at the beat frequency and its first harmonic to attention-related motor processes of an accentuated beat within a metrical grid.

In light of our findings, both the integer ratio bias and the attentional gain may have interacted to lead to these increases. Further research is however needed to disentangle the role of attention during preparation of the motor task from the cognitive processes that enhance the beat based on its simplest binary subdivision (1:2 ratio).

**TABLE S4. Post hoc *t*-tests for Task*Group during the listening to subdivisions and tapping the beat**

| FoI | RHYTHM | GROUP | TASK | | |
|---|---|---|---|---|---|
| | | | **Listening** | **Tapping** | *(Tap - Listen)* |
| 1.25 Hz | Duplets | **Non-Musicians** | M = .016<br>SE = .010 | M = .116<br>SE = .031 | MD = .100<br>*p* = .004 |
| | | **Musicians** | M = .005<br>SE = .009 | M = .194<br>SE = .030 | MD = .189<br>*p* < .001 |
| | | *Mean Diff.*<br>*(M - nonM)* | MD = -.011<br>*p* = .422 | MD = .078<br>*p* = .079 | |
| | Triplets | **Non-Musicians** | M = .007<br>SE = .010 | M = .071<br>SE = .027 | MD = .063<br>*p* = .030 |
| | | **Musicians** | M = .006<br>SE = .009 | M = .232<br>SE = .027 | MD = .226<br>*p* < .001 |
| | | *Mean Diff.*<br>*(M - nonM).* | MD = -.002<br>*p* = .911 | MD = .161<br>*p* < .001 | |
| | Quadruplets | **Non-Musicians** | M = .012<br>SE = .012 | M = .048<br>SE = .023 | MD = .035<br>*p* = .165 |
| | | **Musicians** | M = .023<br>SE = .012 | M = .220<br>SE = .023 | MD = .197<br>*p* < .001 |
| | | *Mean Diff.*<br>*(M - nonM)* | MD = .011<br>*p* = .533 | MD = .172<br>*p* < .001 | |
| | Quintuplets | **Non-Musicians** | M = -.002<br>SE = .007 | M = .022<br>SE = .018 | MD = .024<br>*p* = .194 |
| | | **Musicians** | M = -.003<br>SE = .007 | M = .064<br>SE = .017 | MD = .066<br>*p* = .001 |
| | | *Mean Diff.*<br>*(M - nonM)* | MD = .000<br>*p* = .986 | MD = .042<br>*p* = .096 | |
| 2.5 Hz | Duplets | **Non-Musicians** | M = .119<br>SE = .017 | M = .149<br>SE = .027 | MD = .030<br>*p* = .257 |
| | | **Musicians** | M = .121<br>SE = .016 | M = .302<br>SE = .026 | MD = .181<br>*p* < .001 |
| | | *Mean Diff.*<br>*(M - nonM)* | MD = .002<br>*p* = .923 | MD = .153<br>*p* < .001 | |
| | Triplets | **Non-Musicians** | M = .011<br>SE = .006 | M = .028<br>SE = .019 | MD = .017<br>*p* = .406 |
| | | **Musicians** | M = -.002<br>SE = .006 | M = .152<br>SE = .019 | MD = .155<br>*p* < .001 |
| | | *Mean Diff.*<br>*(M - nonM)* | MD = -.013<br>*p* = .149 | MD = .124<br>*p* < .001 | |

| | | | | | |
|---|---|---|---|---|---|
| (2.5 Hz) | Quadruplets | **Non-Musicians** | M = .014<br>SE = .008 | M = .033<br>SE = .022 | MD = .019<br>$p$ = .382 |
| | | **Musicians** | M = .011<br>SE = .008 | M = .166<br>SE = .021 | MD = .155<br>$p$ < .001 |
| | | *Mean Diff.*<br>*(M - nonM)* | MD = -.002<br>$p$ = .824 | MD = .133<br>$p$ < .001 | |
| | Quintuplets | **Non-Musicians** | M = -.002<br>SE = .006 | M = -.001<br>SE = .009 | MD = .001<br>$p$ = .947 |
| | | **Musicians** | M = .005<br>SE = .005 | M = .023<br>SE = .009 | MD = .019<br>$p$ = .065 |
| | | *Mean Diff.*<br>*(M - nonM)* | MD = .006<br>$p$ = .395 | MD = .024<br>$p$ = .068 | |
| **3.75 Hz** | | **Non-Musicians** | M = .019<br>SE = .003 | M = .029<br>SE = .008 | MD = .010<br>$p$ = .228 |
| | | **Musicians** | M = .015<br>SE = .003 | M = .078<br>SE = .008 | MD = .063<br>$p$ < .001 |
| | | *Mean Diff.*<br>*(M - nonM)* | MD = .003<br>$p$ = .459 | MD = .050<br>$p$ < .001 | |
| **5 Hz** | | **Non-Musicians** | M = .025<br>SE = .003 | M = .028<br>SE = .005 | MD = .003<br>$p$ = .631 |
| | | **Musicians** | M = .032<br>SE = .003 | M = .057<br>SE = .005 | MD = .025<br>$p$ = .005 |
| | | *Mean Diff.*<br>*(M - nonM)* | MD = .006<br>$p$ = .198 | MD = .029<br>$p$ = .029 | |
| **6.25 Hz** | | **Non-Musicians** | M = .022<br>SE = .004 | M = .019<br>SE = .006 | MD = -.003<br>$p$ = .582 |
| | | **Musicians** | M = .022<br>SE = .004 | M = .041<br>SE = .006 | MD = .019<br>$p$ = .001 |
| | | *Mean Diff.*<br>*(M - nonM)* | MD = .000<br>$p$ = .939 | MD = .022<br>$p$ = .011 | |

**Table S4. Post hoc comparisons of the interactions including Task*Group during the listening-to-subdivision tasks.** At each frequency of interest (FoI), neural amplitude means (M) and standard errors (SE) are reported for each Group and Task (and Rhythm when the triple interaction was significant, as in 1.25 and 2.5 Hz). Mean differences (MD) and $p$-values between the Listening and the Tapping tasks are reported in the right column, while mean differences and $p$-values between Musicians and non-Musicians appear below every group division.

**TABLE S5. Post hoc *t*-tests comparing neural entrainment to subdivisions across tasks**

| FoI | RHYTHM | TASK | | |
|---|---|---|---|---|
| | | **Listening** | **Tapping** | *(Tap - Listen)* |
| **3.75 Hz** | Duplets | $M^2$ = -.004<br>SE = .003 | $M^2$ = .050<br>SE = .007 | MD = .054<br>$p$ < .001 |
| | Triplets | $M^3$ = .072<br>SE = .009 | $M^3$ = .111<br>SE = .010 | MD = .038<br>$p$ = .002 |
| | Quadruplets | $M^4$ = -.001<br>SE = .003 | $M^4$ = .046<br>SE = .007 | MD = .047<br>$p$ < .001 |
| | Quintuplets | $M^5$ = .001<br>SE = .003 | $M^5$ = .008<br>SE = .005 | MD = .007<br>$p$ = .219 |
| **6.25 Hz** | Duplets | $M^2$ = .009<br>SE = .007 | $M^2$ = .011<br>SE = .004 | MD = .002<br>$p$ = .792 |
| | Triplets | $M^3$ = .000<br>SE = .003 | $M^3$ = .017<br>SE = .006 | MD = .017<br>$p$ = .001 |
| | Quadruplets | $M^4$ = .000<br>SE = .002 | $M^4$ = .000<br>SE = .002 | MD = .017<br>$p$ = .009 |
| | Quintuplets | $M^5$ = .079<br>SE = .009 | $M^5$ = .075<br>SE = .008 | MD = -.003<br>$p$ = .512 |

**Table S5. Post hoc comparisons of the interaction Task*Rhythm at 3.75 and 6.25 Hz.** Neural amplitude means (M) and standard errors (SE) are reported for each task at each **s**ubdivision rhythm. Mean differences (MD) and *p*-values between the Listening and the Tapping tasks are reported in the right side.

#### TABLE S6. Post hoc *t*-tests comparing neural entrainment to subdivisions across tasks and groups

| FoI | GROUP | TASK: Listening R1 | Listening R2 | Listening *(R1– R2)* | TASK: Tapping R1 | Tapping R2 | Tapping *(R1 – R2)* |
|---|---|---|---|---|---|---|---|
| 1.25 Hz | Non-Musicians | $M^2$ = .016<br>SE = .010 | $M^3$ = .007<br>SE = .010 | MD = .009<br>$p$ = 1 | $M^2$ = .116<br>SE = .031 | $M^3$ = .071<br>SE = .027 | MD = .046<br>$p$ = .280 |
| | | | $M^4$ = .012<br>SE = .012 | MD = .004<br>$p$ = 1 | | $M^4$ = .048<br>SE = .023 | MD = .069<br>$p$ = .005 |
| | | | $M^5$ = -.002<br>SE = .007 | MD = .019<br>$p$ = .950 | | $M^5$ = .022<br>SE = .018 | MD = .095<br>$p$ = .044 |
| | | $M^3$ = .007<br>SE = .010 | $M^4$ = .012<br>SE = .012 | MD = -.005<br>$p$ = 1 | $M^3$ = .071<br>SE = .027 | $M^4$ = .048<br>SE = .023 | MD = .023<br>$p$ = 1 |
| | | | $M^5$ = -.002<br>SE = .007 | MD = .010<br>$p$ = 1 | | $M^5$ = .022<br>SE = .018 | MD = .049<br>$p$ = .581 |
| | | $M^4$ = .012<br>SE = .012 | $M^5$ = -.002<br>SE = .007 | MD = .015<br>$p$ = 1 | $M^4$ = .048<br>SE = .023 | $M^5$ = .022<br>SE = .018 | MD = .026<br>$p$ = 1 |
| | Musicians | $M^2$ = .005<br>SE = .009 | $M^3$ = .006<br>SE = .009 | MD = .000<br>$p$ = 1 | $M^2$ = .194<br>SE = .030 | $M^3$ = .232<br>SE = .027 | MD = -.037<br>$p$ = .539 |
| | | | $M^4$ = .023<br>SE = .012 | MD = -.018<br>$p$ = .931 | | $M^4$ = .220<br>SE = .023 | MD = -.026<br>$p$ = .964 |
| | | | $M^5$ = -.003<br>SE = .007 | MD = .008<br>$p$ = 1 | | $M^5$ = .064<br>SE = .017 | MD = .130<br>$p$ = .002 |
| | | $M^3$ = .006<br>SE = .009 | $M^4$ = .023<br>SE = .012 | MD = -.018<br>$p$ = .747 | $M^3$ = .232<br>SE = .027 | $M^4$ = .220<br>SE = .023 | MD = .012<br>$p$ = 1 |
| | | | $M^5$ = -.003<br>SE = .007 | MD =.008<br>$p$ = 1 | | $M^5$ = .064<br>SE = .017 | MD = .168<br>$p < .001$ |
| | | $M^4$ = .023<br>SE = .012 | $M^5$ = -.003<br>SE = .007 | MD = .026<br>$p$ = .464 | $M^4$ = .220<br>SE = .023 | $M^5$ = .064<br>SE = .017 | MD = .156<br>$p < .001$ |
| 2.5 Hz | Non-Musicians | $M^2$ = .119<br>SE = .017 | $M^3$ = .011<br>SE = .006 | MD = .108<br>$p < .001$ | $M^2$ = .149<br>SE = .027 | $M^3$ = .028<br>SE = .019 | MD = .121<br>$p < .001$ |
| | | | $M^4$ = .014<br>SE = .008 | MD = .105<br>$p < .001$ | | $M^4$ = .033<br>SE = .022 | MD = .116<br>$p < .001$ |
| | | | $M^5$ = -.002<br>SE = .006 | MD = .121<br>$p < .001$ | | $M^5$ = -.001<br>SE = .009 | MD = .150<br>$p < .001$ |
| | | $M^3$ = .011<br>SE = .006 | $M^4$ = .014<br>SE = .008 | MD = -.003<br>$p$ = 1 | $M^3$ = .028<br>SE = .019 | $M^4$ = .033<br>SE = .022 | MD = -.005<br>$p$ = 1 |
| | | | $M^5$ = -.002<br>SE = .006 | MD = .013<br>$p$ = .882 | | $M^5$ = -.001<br>SE = .009 | MD = .029<br>$p$ = .972 |
| | | $M^4$ = .014<br>SE = .008 | $M^5$ = -.002<br>SE = .006 | MD = .016<br>$p$ = .588 | $M^4$ = .033<br>SE = .022 | $M^5$ = -.001<br>SE = .009 | MD = .034<br>$p$ = .766 |
| | Musicians | $M^2$ = .121<br>SE = .016 | $M^3$ = -.002<br>SE = .006 | MD = .124<br>$p < .001$ | $M^2$ = .302<br>SE = .026 | $M^3$ = .152<br>SE = .019 | MD = .150<br>$p < .001$ |
| | | | $M^4$ = .011<br>SE = .008 | MD = .110<br>$p < .001$ | | $M^4$ = .166<br>SE = .021 | MD = .136<br>$p < .001$ |
| | | | $M^5$ = .005<br>SE = .005 | MD = .117<br>$p < .001$ | | $M^5$ = .023<br>SE = .009 | MD = .279<br>$p < .001$ |
| | | $M^3$ = -.002<br>SE = .006 | $M^4$ = .011<br>SE = .008 | MD = -.014<br>$p$ = .532 | $M^3$ = .152<br>SE = .019 | $M^4$ = .166<br>SE = .021 | MD = -.014<br>$p$ = 1 |
| | | | $M^5$ = .005<br>SE = .005 | MD = -.007<br>$p$ = 1 | | $M^5$ = .023<br>SE = .009 | MD = .129<br>$p < .001$ |
| | | $M^4$ = .011<br>SE = .008 | $M^5$ = .005<br>SE = .005 | MD = .007<br>$p$ = 1 | $M^4$ = .166<br>SE = .021 | $M^5$ = .023<br>SE = .009 | MD = .143<br>$p < .001$ |

| | | | | | | | |
|---|---|---|---|---|---|---|---|
| **3.75 Hz** | **All** | $M^2$ = -.004<br>SE = .003 | $M^3$ = .072<br>SE = .009 | MD = -.076<br>$p$ < .001 | $M^2$ = .050<br>SE = .007 | $M^3$ = .111<br>SE = .010 | MD = -.060<br>$p$ < .001 |
| | | | $M^4$ = -.001<br>SE = .003 | MD = -.002<br>$p$ = 1 | | $M^4$ = .046<br>SE = .007 | MD = .004<br>$p$ = 1 |
| | | | $M^5$ = .001<br>SE = .003 | MD = -.004<br>$p$ = 1 | | $M^5$ = .008<br>SE = .005 | MD = .043<br>$p$ < .001 |
| | | $M^3$ = .072<br>SE = .009 | $M^4$ = -.001<br>SE = .003 | MD = .074<br>$p$ < .001 | $M^3$ = .111<br>SE = .010 | $M^4$ = .046<br>SE = .007 | MD = .064<br>$p$ < .001 |
| | | | $M^5$ = .001<br>SE = .003 | MD = .072<br>$p$ < .001 | | $M^5$ = .008<br>SE = .005 | MD = .103<br>$p$ < .001 |
| | | $M^4$ = -.001<br>SE = .003 | $M^5$ = .001<br>SE = .003 | MD = -.002<br>$p$ = 1 | $M^4$ = .046<br>SE = .007 | $M^5$ = .008<br>SE = .005 | MD = -.039<br>$p$ < .001 |
| **5 Hz** | **All** | $M^2$ = .050<br>SE = .004 | $M^3$ = .014<br>SE = .003 | MD = .037<br>$p$ < .001 | No interaction with Task<br>← these values stand for both Listening and Tapping | | |
| | | | $M^4$ = .074<br>SE = .006 | MD = -.024<br>$p$ = .001 | | | |
| | | | $M^5$ = .003<br>SE = .003 | MD = .048<br>$p$ < .001 | | | |
| | | $M^3$ = .014<br>SE = .003 | $M^4$ = .074<br>SE = .006 | MD = -.061<br>$p$ < .001 | | | |
| | | | $M^5$ = .003<br>SE = .003 | MD = .011<br>$p$ = .002 | | | |
| | | $M^4$ = .074<br>SE = .006 | $M^5$ = .003<br>SE = .003 | MD = .072<br>$p$ < .001 | | | |
| **6.25 Hz** | **All** | $M^2$ = .009<br>SE = .007 | $M^3$ = .000<br>SE = .003 | MD = .009<br>$p$ = 1 | $M^2$ = .011<br>SE = .004 | $M^3$ = .017<br>SE = .006 | MD = -.006<br>$p$ = .623 |
| | | | $M^4$ = .000<br>SE = .002 | MD = .009<br>$p$ = 1 | | $M^4$ = .000<br>SE = .002 | MD = -.006<br>$p$ = .503 |
| | | | $M^5$ = .079<br>SE = .009 | MD = -.070<br>$p$ < .001 | | $M^5$ = .075<br>SE = .008 | MD = .064<br>$p$ < .001 |
| | | $M^3$ = .000<br>SE = .003 | $M^4$ = .000<br>SE = .002 | MD = .000<br>$p$ = 1 | $M^3$ = .017<br>SE = .006 | $M^4$ = .000<br>SE = .002 | MD = .000<br>$p$ = 1 |
| | | | $M^5$ = .079<br>SE = .009 | MD = -.078<br>$p$ < .001 | | $M^5$ = .075<br>SE = .008 | MD = -.058<br>$p$ < .001 |
| | | $M^4$ = .000<br>SE = .002 | $M^5$ = .079<br>SE = .009 | MD = -.079<br>$p$ < .001 | $M^4$ = .000<br>SE = .002 | $M^5$ = .075<br>SE = .008 | MD = -.058<br>$p$ < .001 |

**Table S6. Post hoc comparisons of the significant main effects and interactions related to the subdivision rhythms.** At each frequency of interest, the table reports the neural amplitude means (M) and standard errors (SE) for both the Listening and Tapping tasks (except at 5Hz, where there was no interaction *Rhythm***Task*). $M^2$ stands for duplets, $M^3$ for triplets, $M^4$ for quadruplets, and $M^5$ for quintuplets. The table reports mean differences (MD) between the mean under R1 and the mean under R2, as well as their corresponding $p$-values on the right side of every task. The values for musicians and non-musicians are only shown at 1.25 and 2.5Hz, as there was a significant triple interaction involving the between factor *Group*. These post hoc comparisons provide further insights into the effects of subdivision and music training on synchronized neural amplitudes.

### 5. ADDITIONAL NON-PARAMETRIC TESTS FOR TAPPING CONSISTENCY

**Table S7. Nonparametric ANOVA and post hoc tests for musicians' tapping consistency**

*Friedman Test*

| Factor | $\chi^2_F$ | df | p | Kendall's W |
|---|---|---|---|---|
| Subdivision | 29.47 | 3 | < .001 | 0.614 |

**Conover Test**

*Conover's Post Hoc Comparisons - Subdivision*

| | | T-Stat | df | $W_i$ | $W_j$ | $r_{rb}$ | p | $p_{bonf}$ | $p_{holm}$ |
|---|---|---|---|---|---|---|---|---|---|
| duplet | triplet | 0.640 | 45 | 48.00 | 51.00 | 0.132 | .525 | 1.000 | 1.000 |
| | quadruplet | 0.640 | 45 | 48.00 | 45.00 | 0.529 | .525 | 1.000 | 1.000 |
| | quintuplet | 6.829 | 45 | 48.00 | 16.00 | 1.000 | < .001 | < .001 | < .001 |
| triplet | quadruplet | 1.280 | 45 | 51.00 | 45.00 | 0.412 | .207 | 1.000 | .621 |
| | quintuplet | 7.470 | 45 | 51.00 | 16.00 | 1.000 | < .001 | < .001 | < .001 |
| quadruplet | quintuplet | 6.189 | 45 | 45.00 | 16.00 | 1.000 | < .001 | < .001 | < .001 |

*Note.* Grouped by subject.

*Note.* Rank-biserial correlation based on individual signed-rank tests.

**Table S8. Nonparametric ANOVA and post hoc tests for non-musicians' tapping consistency**

*Friedman Test*

| Factor | $X^2_F$ | df | p | Kendall's W |
|---|---|---|---|---|
| Subdivision | 26.60 | 3 | < .001 | 0.591 |

**Conover Test**

*Conover's Post Hoc Comparisons - Subdivision*

| | | T-Stat | df | $W_i$ | $W_j$ | $r_{rb}$ | p | $p_{bonf}$ | $p_{holm}$ |
|---|---|---|---|---|---|---|---|---|---|
| duplet | triplet | 3.632 | 42 | 54.00 | 37.00 | 0.750 | < .001 | .005 | .002 |
| | quadruplet | 2.778 | 42 | 54.00 | 41.00 | 0.800 | .008 | .049 | .016 |
| | quintuplet | 7.692 | 42 | 54.00 | 18.00 | 0.983 | < .001 | < .001 | < .001 |
| triplet | quadruplet | 0.855 | 42 | 37.00 | 41.00 | -0.017 | .398 | 1.000 | .398 |
| | quintuplet | 4.060 | 42 | 37.00 | 18.00 | 0.900 | < .001 | .001 | < .001 |
| quadruplet | quintuplet | 4.914 | 42 | 41.00 | 18.00 | 1.000 | < .001 | < .001 | < .001 |

*Note.* Grouped by subject.

*Note.* Rank-biserial correlation based on individual signed-rank tests.
